# Circadian timing does not modulate human temporal contrast sensitivity

**DOI:** 10.1101/2025.11.10.687546

**Authors:** Hannah Sophie Heinrichs, Manuel Spitschan

**Affiliations:** Max Planck Institute for Biological Cybernetics, Tübingen, Germany; Technical University of Munich, Munich, Germany; Technical University of Munich, TUM Institute for Advanced Study (TUM-IAS), Germany TUMCREATE, Singapore

**Keywords:** Temporal contrast sensitivity, Silent substitution, Circadian rhythm, Repeated-measures design, Cones

## Abstract

**Introduction:** Previous studies have reported diurnal variations in colour perception and luminance contrast sensitivity, suggesting modulation of visual processing by the circadian pacemaker. To clarify potential time-of-day effects, we examined whether cone-mediated temporal contrast sensitivity is modulated by circadian timing.

**Methods:** We disentangled circadian and homeostatic influences on human visual performance using a 40-hour forced desynchrony protocol. Twelve participants (5 women, 24.4 ± 2.9 years, range 19–30) followed a repeating 3.75 h cycle comprising 2 h 30 min of wakefulness and 1 h 15 min of sleep under constant environmental conditions. Circadian phase was determined from 46 salivary melatonin samples collected per participant. During wake periods, participants repeatedly completed temporal contrast sensitivity tasks at 2 Hz and 8 Hz, probing luminance (L+M+S) and chromatic (L–M, S-cone-directed) modulations. We tested for circadian modulation of visual performance by comparing multilevel cosinor and uninformative Bayesian models.

**Results:** All participants showed robust melatonin rhythms (mean *τ* = 24.18 ± 0.39 h). Bayesian inference provided decisive evidence against circadian variation in luminance or chromatic contrast sensitivity.

**Discussion:** Within this controlled protocol, we found no detectable near-24-hour modulation of cone-mediated temporal contrast sensitivity. Previously reported time-of-day effects may instead reflect homeostatic influences, pupil-linked changes in retinal illuminance, or mechanisms outside the cone-mediated pathways probed here.

**Significance:** By separating circadian phase from sleep–wake-related influences, this study clarifies when circadian modulation of visual performance can, and cannot, be inferred.

## Introduction

Following the discovery of the melanopsin photoreceptor system (Berson, Dunn, & Takao, 2002; Hattar, Liao, Takao, Berson, & Yau, 2002), research on the effects of light on circadian rhythms and human physiology has expanded rapidly (Vetter et al., 2021). Yet, little attention has been paid to the reverse relationship, namely how light signals and downstream (non-)image-forming functions are endogenously modulated by the circadian rhythm. Such modulation may occur at multiple levels along the visual pathway, from the retina to higher cortical areas. The dynamic nature of visual processing, including its temporal variability, suggests an interdependent relationship between sensory systems and biological timekeeping (Andrade, 2022), warranting a systematic investigation of both directions of this potentially bidirectional link.

### Photoreceptors and post-receptoral mechanisms

At the retina, light is absorbed by three classes of photoreceptors: rods, cones, and intrinsically photosensitive retinal ganglion cells (ipRGCs). Rods support vision in low light, while cones mediate colour vision and have the highest density in the fovea. Cones comprise three classes with overlapping spectral sensitivities: short-, medium-, and long-wavelength-sensitive, or S, M, and L cones. Their peak sensitivities lie at approximately 440 nm, 530 nm, and 560 nm, respectively. Relative activation across these cone classes enables trichromatic colour vision. Because individual photoreceptors do not distinguish between wavelength and intensity, colour perception depends on comparing the relative activity across multiple cone types. In other words, the response of any single cone is ambiguous: a given response could be produced either by changing intensity at one wavelength or by presenting a different wavelength at another intensity (principle of univariance; (Rushton, 1972)). Photoreceptors convert optical radiation into electrical signals that are processed within and across multiple retinal layers before being relayed to subcortical and cortical structures. Signals from L, M, and S cones are integrated and recombined through post-receptoral mechanisms, enabling discrimination of luminance contrast and colour along distinct axes. Luminance contrast is driven by combined cone input (L+M+S), while colour vision is mediated by two opponent channels: L–M (the red–green mechanism) and S–[L+M] (the blue–yellow mechanism) (Stockman & Brainard, 2010).

### Circadian control in retinal function

Many physiological and psychological functions, including retinal sensitivity, follow circadian variation governed by endogenous biological clocks. Studies in rodents indicate that photoreceptor signalling and retinal tissue exhibit diurnal or circadian modulation (Tosini, Pozdeyev, Sakamoto, & Iuvone, 2008; Danilenko, Plisov, Cooper, Wirz-Justice, & Hébert, 2011). Within the photoreceptor layer, clock genes show rhythmic expression (Liu, Zhang, & Ribelayga, 2012), and electroretinography (ERG) reveals circadian rhythmicity in retinal function (Cameron & Lucas, 2009; Hankins, Jones, & Ruddock, 1998). These rhythms appear to be generated by peripheral clocks within the retina, although they may also be modulated by the central circadian pacemaker in the suprachiasmatic nucleus (SCN) via neuronal or humoral pathways (Roenneberg & Merrow, 2016). The interplay between local and central mechanisms highlights the complexity of circadian regulation in the visual system.

### Diurnal variability in visual performance

While circadian rhythms in retinal physiology are established, evidence for their influence on visual performance is limited. Human studies report diurnal variation in luminance and colour sensitivity (Andrade, Neto, Oliveira, Santana, & Santos, 2018; Andrade, Cristino, Santos, Oliveira, & Santos, 2018; Andrade, Rezende, Figueiredo, Farias, & Santos, 2019; Tassi, Pellerin, Moessinger, Eschenlauer, & Muzet, 2000; Tassi, Pellerin, & Muzet, 1999; Tassi & Pins, 1997; O’Keefe & Baker, 1987; Morita, Ohyama, & Tokura, 1994). However, these findings are inconsistent and difficult to compare because of methodological differences. Many studies on luminance perception focused on spatiotemporal frequency rather than circadian rhythmicity, lacking adequate sampling across the full 24-hour cycle. For example, Andrade, Cristino, et al. (2018) found daily maxima in visual sensitivity along confusion axes that contrast with morning–evening differences in cone pathway responsivity observed in ERG recordings by Danilenko et al. (2011). The underlying mechanisms of these fluctuations – whether homeostatic, circadian, or both – remain unclear. Existing studies also tend to use stimuli defined in non-cone colour spaces or monochromatic light (Morita et al., 1994; Andrade, Cristino, et al., 2018), which limits precision in linking variability to specific photoreceptive or post-receptoral mechanisms. Only targeted modulation of defined pathways could reveal whether central circadian clocks influence perceptual outcomes such as colour discrimination and luminance contrast perception. Moreover, most studies test participants under naturalistic sleep–wake schedules, preventing separation of endogenous circadian from behavioural or homeostatic influences.

### Circadian rhythm and sleep homeostasis

Although retinal clocks are established, the interaction between local processes, the central pacemaker, and homeostatic regulation remains unresolved. Visual performance depends on psychological and physiological states – motivation, vigilance, sleepiness, and metabolic factors (Barlow, Farell, & Khan, 2003) – which fluctuate on similar timescales as the circadian rhythm. The body alternates between sleep and wakefulness in a roughly 24-hour cycle, regulating rest–activity and feeding–fasting rhythms. According to the two-process model, circadian and homeostatic processes are independent but interacting regulators of physiological state. To attribute diurnal variations in visual outcomes to circadian modulation, their effects must be disentangled from sleep–wake–related influences. Two experimental paradigms enable such separation: the constant routine (CR) and forced desynchrony (FD) protocols (Wirz-Justice, 2007). In a CR, participants remain awake for about 40 hours under dim light, constant posture, and controlled temperature and nutrition, allowing assessment of circadian effects while accounting for sleep deprivation (Kräuchi, 2007; Cajochen, Knoblauch, Kräuchi, Renz, & Wirz-Justice, 2001). FD protocols impose non-24-hour cycles (e.g., 20- or 28-hour “days”), preventing circadian entrainment while maintaining regular sleep opportunities (Wang et al., 2022). This approach enables estimation of independent circadian and homeostatic components. Although FD protocols are resource-intensive, shorter variants (multiple-nap protocols) maintain low, stable sleep pressure through frequent naps (Cajochen et al., 2001; Wirz-Justice, 2007). To date, no FD or CR study has systematically examined visual performance outcomes (Wang et al., 2022, see Supplementary Information).

### Open research questions

In the current body of literature, significant gaps remain in the understanding of the dynamics between the visual and the circadian system with regard to visual functions. Specifically, it is unclear whether and how early sensory rhythmicity translates into functionally relevant performance outcomes, and what biological mechanisms drive changes in visual performance measures.

### Aim of the study

To address these questions, we designed a study combining knowledge from vision science and chronobiology to investigate whether post-receptoral mechanisms underlying colour and luminance contrast sensitivity are subject to circadian rhythms. We used a shortened forced desynchrony (FD) protocol with ultra-short sleep–wake cycles in a constant environment to minimise sleep pressure and control environmental and behavioural influences.

### Hypotheses

We propose that if circadian modulation acts at an early sensory stage, it may be detectable as variation in visual performance across an individual’s circadian period. We therefore test whether colour and luminance contrast sensitivity show circadian modulation. Specifically, we examine three post-receptoral directions: L+M+S (achromatic luminance modulation), L–M (nominal red-green cone-opponent modulation), and S-cone-directed modulation with L-and M-cone activation held constant. These stimulus directions were chosen because they map onto canonical post-receptoral pathways relevant to luminance contrast and colour discrimination (Engel, Zhang, & Wandell, 1997). Given the differing temporal contrast sensitivity (TCS) functions for luminance and chromatic pathways (Gelfand & Horwitz, 2018), we probe each direction at two temporal frequencies. A lower frequency (2 Hz) was selected to weight the measurement toward chromatic sensitivity, whereas a higher frequency (8 Hz) was selected to weight the measurement relatively more toward luminance sensitivity. We do not interpret either frequency as exclusively isolating one mechanism; rather, the two frequencies provide complementary sampling points on the known temporal tuning functions.

This approach is exploratory, but it allows us to ask whether any circadian modulation differs across stimulus directions and across portions of the temporal response functions where luminance and chromatic mechanisms are expected to contribute differently.

These considerations lead us to six specific hypotheses, corresponding to the three post-receptoral mechanisms tested at each frequency. We test circadian modulations in TCS in different models:

1. “L+M+S (luminance), 2 Hz”,
2. “L+M+S (luminance), 8 Hz”,
3. “L–M (red-green), 2 Hz”,
4. “L–M (red-green), 8 Hz”,
5. “S (blue-yellow), 2 Hz”, and
6. “S (blue-yellow), 8 Hz” .

## Methods

The study design and experimental protocol used in the present work were previously described by Heinrichs and Spitschan (2025). It focuses on visual functions but also includes a range of other outcomes related to ocular physiology, cardio-metabolic dynamics, and indicators of cognitive functioning and psychological state.

Participants initially enrolled in a study comprising periods of in-field circadian stabilisation and multiple in-laboratory experimental sessions. The protocol began with a seven-day stabilisation period, followed by a training session, another seven-day stabilisation period, and a dim-light evening session. This was followed by a 40-hour circadian protocol adhering to an ultra-short sleep–wake rhythm. After the forced desynchrony protocol, participants re-stabilised to their agreed sleep–wake schedule for an additional five days. During the wake periods of the 40-hour forced desynchrony, participants completed psychophysical experiments assessing temporal contrast sensitivity. All data were collected between January and October 2024.

### Participant recruitment

Recruitment methods included flyers placed in public areas, online advertisements, and social media outreach. The flyers directed interested individuals to a study website providing detailed information and access to the online screening. The online screening consisted of seven questionnaires completed sequentially. Individuals with normal or corrected-to-normal vision and regular sleep patterns were eligible, as outlined in Tables 1 and 2. A total of 13 participants were deemed eligible for laboratory screening, and 12 completed the experiment with valid data. One participant’s data were excluded due to a technical issue: during their experimental session, display settings were inadvertently changed, resulting in inconsistent colour presentation that compromised data validity. Due to resource and time constraints, recruitment was concluded once the maximum target sample size was reached. The study was approved by the Ethics Committee of the Technical University of Munich (2023-369-S-SB). All participants provided written informed consent. All research was conducted in accordance with the Declaration of Helsinki.

**Table 1:** Inclusion criteria.

| Domain | Assessment method | Criterion | Time of Screening |
| --- | --- | --- | --- |
| Age | Self-report | $\geq 18$ years<br>$\leq 35$ years | Online screening |
| Physical health | Self-report | Good physical health | Online screening |
| Mental health | Self-report | Good mental health | Online screening |
| Ocular health | Ophthalmological examination | Good ocular health | In-person screening |
| Visual acuity | Landolt C | Normal or corrected-to-normal vision | In-person screening |
| Colour vision | HRR Pseudoisochromatic Plates (4th edition) | Normal colour vision | In-person screening |
| Sleep and circadian regularity | Self-report | Preference of high ranking participants | Online screening |
| Hormone status | Self-report | Women preferably taking oral contraception, Men without any hormone treatments | Online screening |

**Table 2:**
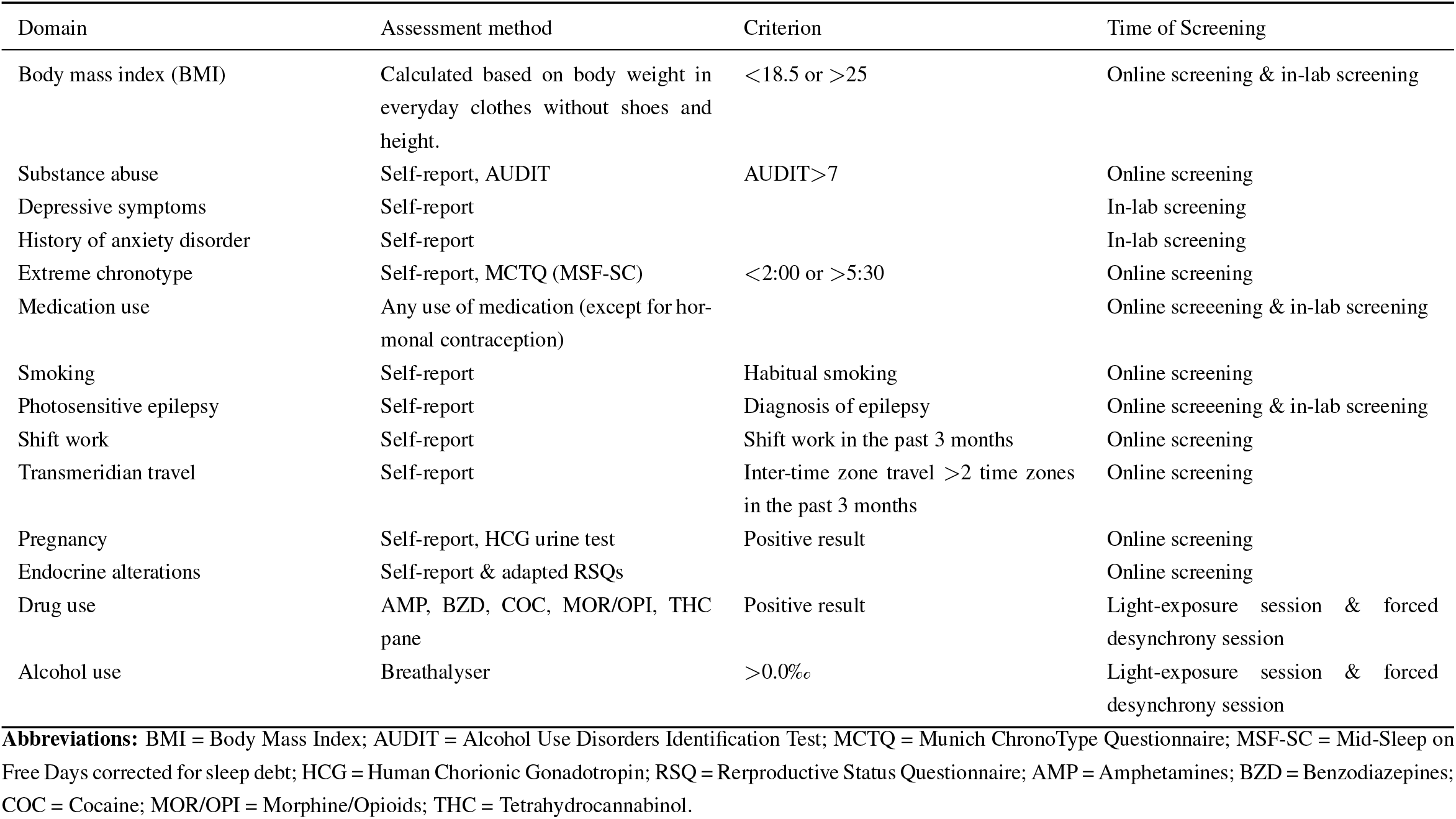
Exclusion criteria.

### Participant characteristics

The final study sample consisted of healthy volunteers (*N* = 12; mean age = 24.42 ± 2.87 years, range = [19–30]). The sample included five females, two of whom used oral contraceptives, while the remaining three had natural menstrual cycles without hormonal influence. One participant had corrected vision and wore glasses during the experimental session, removing them during visual testing. Participants reported regular sleep patterns prior to the experimental session (Table 3). During an in-person screening visit, participants toured the facilities and had the opportunity to ask questions. Physical health was confirmed through measurement of blood pressure, VO_2_max, body temperature, and the absence of psychological or medical conditions that could compromise the study protocol or confound outcome measures. Immediately before the experimental session, participants were tested for abstinence from alcohol and drugs using a breathalyser (ACE GmbH, Freilassing, Germany) and a urine drug test (nal von minden GmbH, Moers, Germany). These tests confirmed compliance with study conditions regarding substance intake for all participants.

**Table 3:** Participants’ sleep and sleep-related characteristics.

| *Measure (Abbreviation)* | *Mean ± St.d.* | *Range* | *Reference and description* |
| --- | --- | --- | --- |
| Mid-sleep on free days corrected for sleep debt (MSF<sub>SC</sub>) [hh:mm] | 4:20±0:53 | [2:34; 5:30] | Roenneberg et al. (2003); collected using the μMCTQ to estimate midpoint of sleep on free days, adjusted for sleep debt. |
| Pittsburgh Sleep Quality Index (PSQI) | 2.27±1.62 | [0.0; 5.0] | Buysse et al. (1989); assesses subjective sleep quality over a 1-month interval. |
| Epworth Sleepiness Scale (ESS) | 4.27±1.79 | [1.0; 7.0] | Johns (1991); measures habitual daytime sleepiness via self-reported questionnaire. |
| Sleep Regularity Questionnaire (SRQ) | 32.82±4.42 | [22.0; 38.0] | Dzierzewski et al. (2021); evaluates sleep regularity based on timing and variability in sleep patterns. |

### Study design

For psychophysical testing, we used a series of two-alternative forced-choice (2AFC) tasks to repeatedly assess the temporal contrast sensitivity of three distinct retinal mechanisms. Testing was performed multiple times during wake periods across a 40-hour interval, covering approximately one and a half circadian cycles. Figure 1 provides a schematic overview of the study protocol, psychophysical procedure, experimental design, and stimulus design. During the forced desynchrony protocol, various assessments targeting visual and non-visual functions were conducted. A detailed description of the procedures performed during wake periods (e.g., pupillometric tests, vigilance assessments, ocular measurements, questionnaires, and subjective ratings) and of continuously monitored outcomes (e.g., skin temperature, core body temperature, and ECG) is available in the published protocol (Heinrichs & Spitschan, 2025). Other outcome variables collected during the study will be analysed separately, as their corresponding hypotheses and analyses are independent of the present results. These data will be published elsewhere.

**Figure 1.**
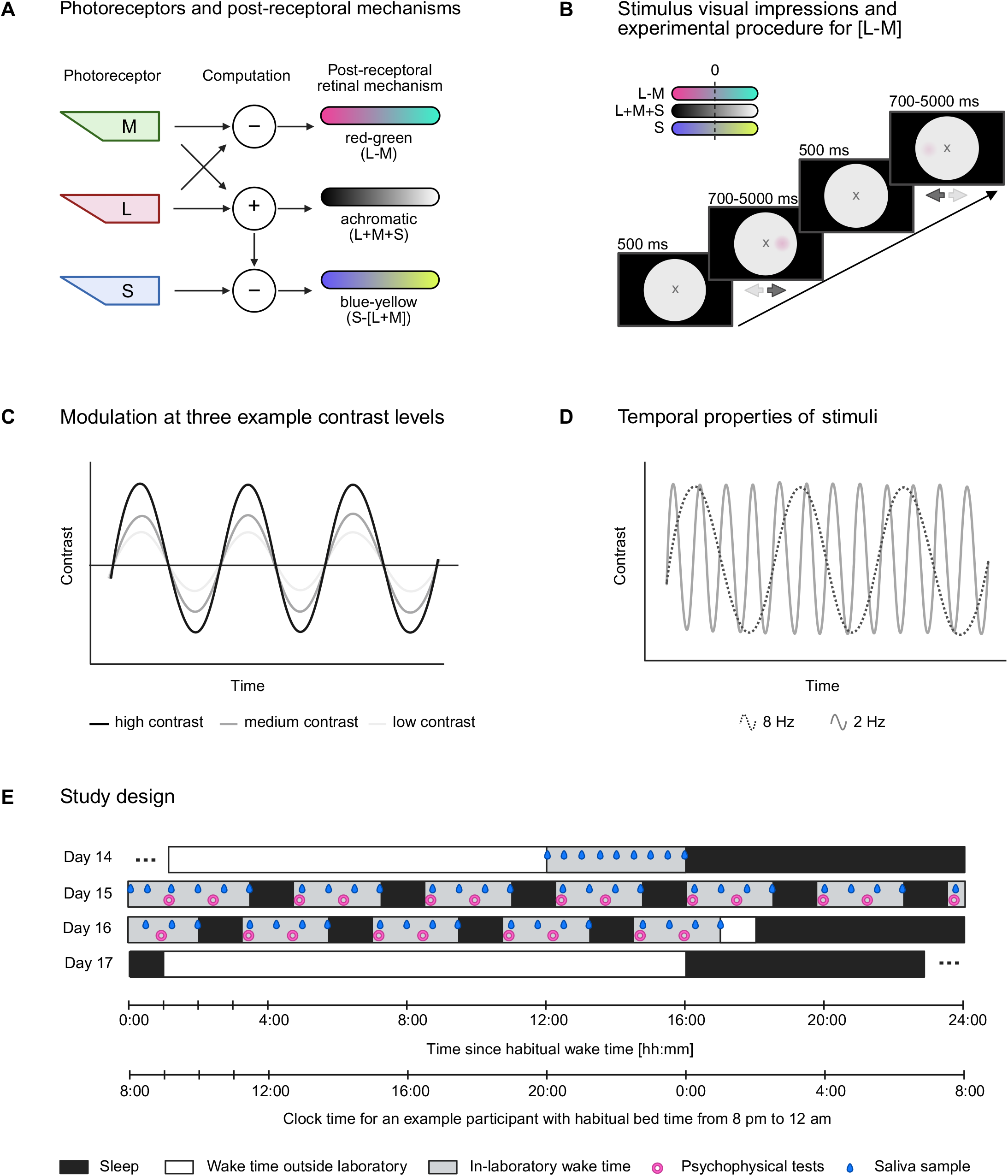
(Previous page.) Schematic overview of the study protocol, psychophysical procedure, experimental design, and stimulus design. (A) Computational framework motivating the stimulus design: cone signals are recombined by post-receptoral pathways relevant to luminance contrast and colour vision. (B) The psychophysical experiment comprised three stimulus directions: L+M+S modulation, L–M modulation, and S-cone-directed modulation with L- and M-cone activation held constant. Approximate visual impressions are shown schematically. Stimuli were presented on a monitor viewed through a circular cutout. (C) The task used an adaptive staircase to adjust stimulus contrast. (D) Each task contained two temporal frequencies, 2 Hz and 8 Hz. (E) Overview of the 40-hour laboratory protocol, including psychophysical tests and saliva sampling.

#### Circadian study protocol

We used a 40-hour forced desynchrony (FD) protocol to decouple participants’ sleep–wake rhythm from their intrinsic circadian rhythm. The protocol consisted of 11 experimental blocks, interspersed with sleep opportunities, in a repeating 3.75-hour cycle comprising 2 hours 30 minutes of wakefulness and 1 hour 15 minutes of sleep (sleep-wake ratio or 1:2). Since the first block took place after an 8-hour adaptation night, the protocol began with a wake period. In the 11th block, participants were sent home after the final wake period. Prior to the experimental session, participants underwent a 7-day stabilisation period. During this time, they maintained a self-selected sleep–wake rhythm with consistent bed and wake times within ±30 minutes. They also adhered to a self-selected meal schedule of either two, three, or four meals per day. Compliance with the stabilisation protocol was monitored using REDCap online questionnaires to record sleep and meal logs, as well as activity tracking devices. Only participants who fully complied with these requirements were permitted to proceed to the experimental session. There was one case of non-compliance, in which the experimenter decided to reschedule the session. In the evening before the FD protocol began, participants entered the laboratory and spent 4 hours in a dim-light environment. The purpose of this period was to standardise light exposure history for at least 12 hours before the start of the experiment. Participants then received a full night of 8-hour sleep to acclimatise to the controlled laboratory environment. The FD protocol commenced the following morning, 1 hour after each participant’s habitual wake time. During this transitional hour, participants were equipped with all necessary devices for the experiment.

#### Saliva collection

Throughout the study, hormone data were obtained from saliva samples to characterise melatonin and cortisol profiles across the circadian rhythm for subsequent analyses. Saliva samples (at least 1 mL) were collected approximately every 45 minutes during the FD protocol and twice during the hour between wake-up and protocol start, using Salivettes^®^ (Sarstedt, Nürnberg, Germany). Participants chewed on a cotton swab for 3 minutes and then placed it into a plastic collection tube. Samples were centrifuged for 3 minutes at 3,000 rpm and stored at –20°C for temporary preservation. Biological samples were first stored in the laboratory and subsequently sent to external facilities for processing. Melatonin concentrations were determined by radioimmunoassay (RIA, RK-DSM2, NovoLytiX GmbH, Switzerland; typical specifications: limit of quantification 0.5–50 pg/mL, detection limit 0.2 pg/mL, mean intra-assay precision 7.9

### Experimental setup

Psychophysical tests were presented using the Metropsis system (Cambridge Research Systems, Rochester, UK) on a colour-calibrated, three-primary 32-inch LCD monitor with 10-bit colour resolution and a 120 Hz refresh rate. The monitor was factory-calibrated by the manufacturer during device setup. The field of view was restricted by a circular blackout cutout, approximately 38 cm (15 inches) in diameter, placed in front of the monitor.

#### Pupillary relay system

All psychophysical testing was conducted with a retinal relay system. Participants viewed the stimulus monocularly with their dominant eye through an entry pupil positioned 1.5 metres from the participant. The image at the entry pupil plane was projected onto the human pupil using a multi-lens optical system. The entry pupil consisted of a fixed aperture of 1.5 mm, ensuring that retinal irradiance remained constant and unaffected by light-induced pupil constriction. The experimental setup was located in a designated area enclosed by light-blocking curtains to minimise stray light.

#### Experimental procedure

At the onset of the FD protocol, refractive correction was adjusted once for each participant using a standardised subjective focusing procedure with a Snellen chart viewed through the optical system. No further corrections or adjustments were made during the protocol. The protocol consisted of 11 experimental blocks, during which participants repeatedly completed three psychophysical staircase tasks to assess the temporal contrast sensitivity of three post-receptoral retinal mechanisms (L+M+S, L–M, and S). All tests were delivered monocularly to the participant’s dominant eye, determined in a prior screening session using the Miles test (Miles, 1930). In this test, participants fully extended their arms and created a triangular opening between their thumbs and index fingers, focusing on a distant object through the opening with both eyes. They alternately closed each eye and reported which eye kept the object in view. The eye that maintained the object when the other was closed was identified as the dominant eye. The dominant eye was selected for testing because prolonged visual attention with the non-dominant eye may induce fatigue. Each trial was a two-alternative forced-choice (2AFC) location-discrimination judgment. A stimulus appeared randomly on either the left or right side of the display, at 2° visual angle from fixation, and participants indicated the side on which the modulation occurred. The two stimulus locations were treated as equivalent within observers; any systematic left-right sensitivity difference would therefore contribute noise rather than a circadian signal. Within a given test, one post-receptoral direction was probed and the two temporal frequencies (2 and 8 Hz) were interleaved. Stimulus contrast was adjusted adaptively with a two-up-one-down staircase based on the sequence of 2AFC responses. Here, “contrast” refers to the nominal modulation depth of the targeted stimulus direction relative to the grey background. Before the first reversal, two consecutive correct responses reduced the presented contrast by 50%; after the first reversal, the decrease step was 12.5%. A single incorrect response increased the presented contrast by 25%. Catch trials (10%) were included, yielding an average target performance level of 79.1%. Maximum contrast levels were set to 30% for the L+M+S direction, 51% for the S direction, and 6% for the L–M direction due to the display’s gamut limits for those stimulus directions. Starting contrasts were set to 15% for the L+M+S direction, 51% for the S direction, and 6% for the L-M direction. Each frequency-specific staircase terminated after six reversals. The contrast threshold estimate was calculated as the mean of the final four peaks and troughs of the staircase, and the associated uncertainty was derived from the same set of reversal values (Figure 3).

**Figure 2.**
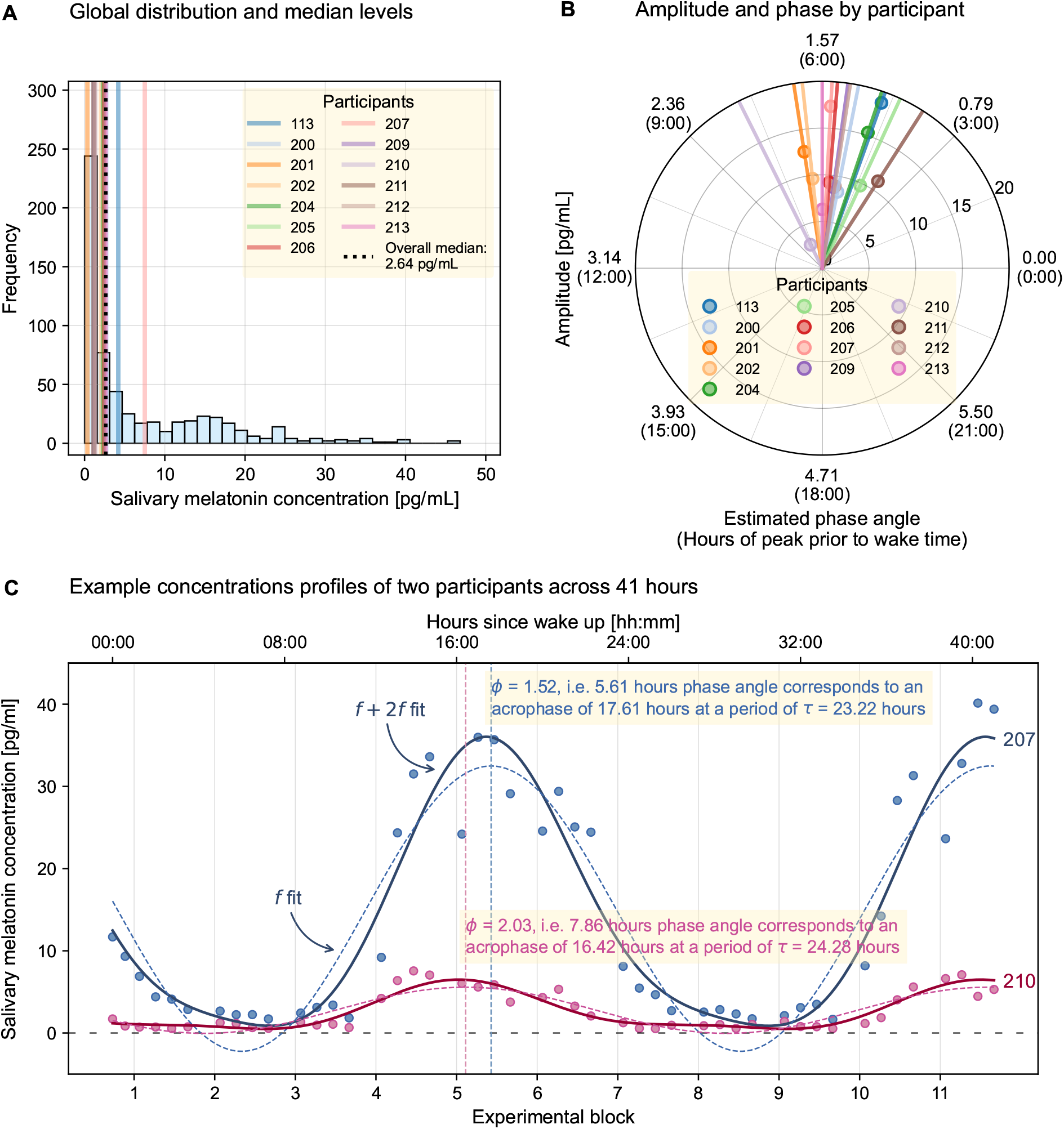
Overview of data and statistics illustrating periodic melatonin secretion. Melatonin sampling started after wake-up, 1 hour before the start of the 40-hour forced desynchrony protocol, covering a total of 41 hours. (A) Distribution of salivary melatonin concentrations across all samples, illustrating a strong accumulation of near-zero values. Coloured lines indicate participant-specific median concentrations, most being zero or near zero; the overall median (2.64 pg/mL) is shown by the black dashed line. (B) Individual amplitude and phase estimates from the fundamental component (*f*) of a bi-harmonic (*f* + 2*f*) cosinor model. Amplitude is represented by the radial distance from the origin, and phase (acrophase) by the angular position on the interval [0, 2*π*[, corresponding to 0:00–24:00 hours advance relative to wake time. Anchor of phase = 0 corresponds to an acrophase of 0:00; positive values, e.g. 1.57, indicate peak of cosinor 6:00 hours before wake time. (C) Example melatonin profiles of two participants (207 and 210) across 41 hours. Dots represent measured melatonin concentrations; solid lines show the fitted cosinor model including the fundamental and second harmonic (*f* + 2*f*); dashed lines indicate the fundamental component alone, which was obtained from the bi-harmonic model, and from which amplitude and phase in subplot B were derived. Dashed vertical lines mark the acrophase (peak) of the fundamental component, corresponding to the phase angles shown in panel B. For visual clarity, the acrophase is plotted one cycle (approx. 24 hours) later so that it is marked after wake time.

**Figure 3.**
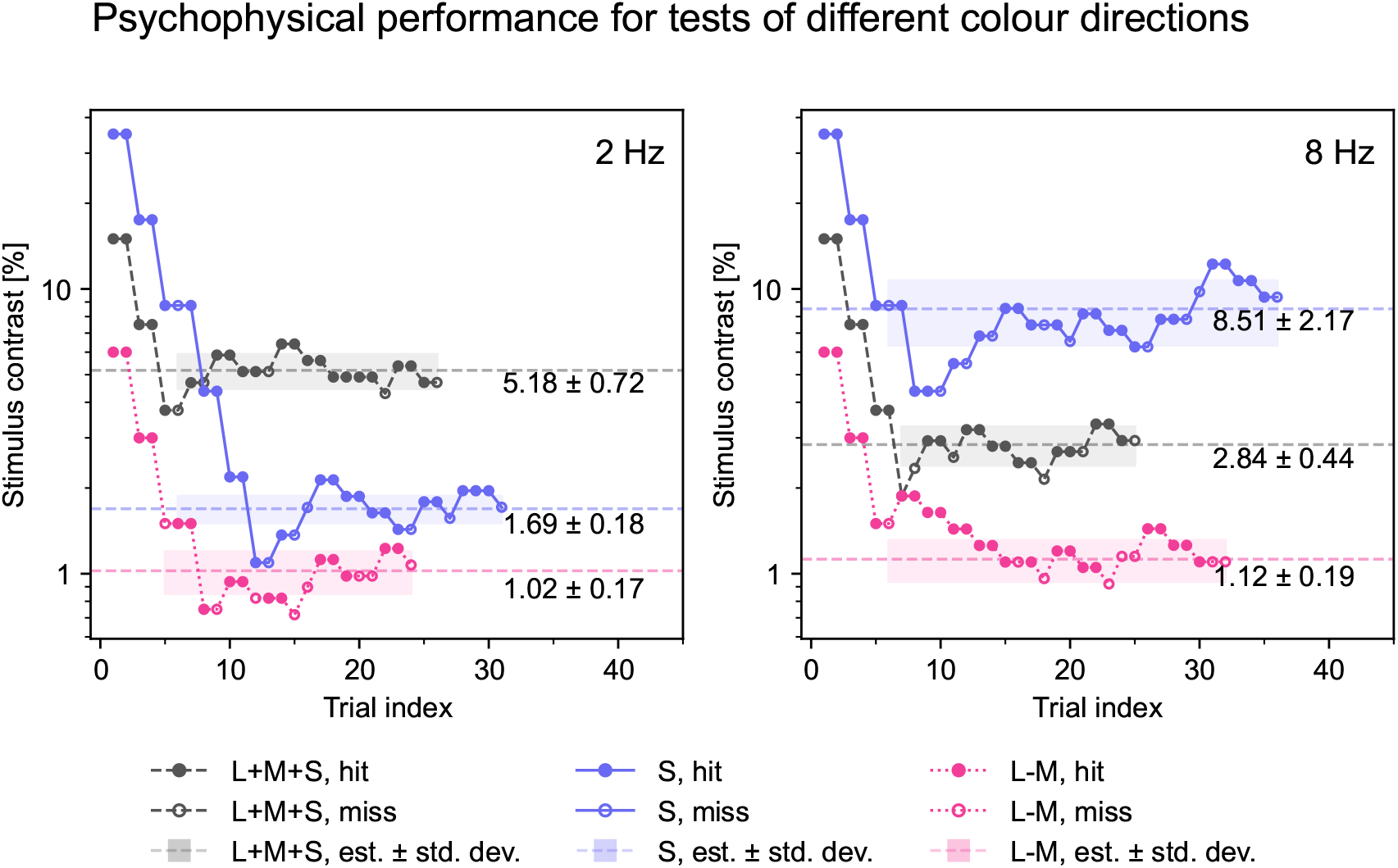
Example staircase data for location discrimination in three tests targeting the retinal mechanisms “L+M+S”, “L–M”, and “S”. The data are taken from three example tests (each containing both frequencies) from participant 212 in the third experimental block. Frequency-specific staircases are shown separately, with 2 Hz on the left and 8 Hz on the right. Contrast modulation was based on Weber contrast calculations, therefore stimulus contrast is expressed in percent. Contrast threshold estimate per condition and standard deviation is provided in the figure. All following analysis were done on contrast sensitivity which is defined at inverse of the final stimulus contrast threshold. Each point indicates the presented stimulus contrast on one trial; filled points are correct responses (hits) and open points are incorrect responses (misses). The dashed line indicates the final threshold estimate, and the shaded band indicates the standard deviation of the reversal values used to estimate that threshold.

#### Silent substitution stimuli at different frequencies

The tasks modulated three nominal post-receptoral directions against a grey background with a luminance of 59.88 cd/m^2^ with a chromaticity of *x, y* = (0.2997, 0.3283): luminance contrast (L+M+S), red-green cone-opponent modulation (L–M, with S-cone activation held constant), and S-cone-directed modulation (with L- and M-cone activation held constant). Stimuli for luminance contrast could theoretically reach a luminance of 77.84 cd/m^2^ at +30% Weber contrast for the L+M+S, while chromatic stimuli are inherently isoluminant with the background. Because the modulation was symmetric about the background, the mean luminance over time was 59.88 cd/m^2^. The silent substitution method is designed to modulate one or more photoreceptor classes while holding others constant (Spitschan & Woelders, 2018). Unlike monochromatic stimulation, which necessarily co-modulates multiple photoreceptor classes because of overlapping spectral sensitivities, silent substitution uses the spectral output of the display primaries together with assumed observer spectral sensitivities to target specific directions in cone-excitation space. To generate stimuli, standard-observer cone fundamentals (CIE, 2006) were supplied to the software, which used linear inversion to convert LMS values into device RGB coordinates. We did not perform observer-specific calibrations of L:M ratio, prereceptoral filtering, or macular pigment density; the stimuli should therefore be interpreted as nominally cone-directed under standard-observer assumptions rather than as perfectly mechanism-pure in every participant. Stimuli consisted of spatially homogeneous, temporally modulated circular fields with a Gaussian spatial envelope to ensure smooth blending into the grey background. Contrast modulation followed a sinusoidal function over time, with the midpoint corresponding to the background luminance.

#### Stimulus frequency selection

We tested each stimulus direction at two temporal frequencies, 2 Hz and 8 Hz. These frequencies were chosen to sample different portions of the known temporal response functions of chromatic and luminance mechanisms rather than to exclude one mechanism entirely. At 2 Hz, chromatic sensitivity is typically relatively stronger; at 8 Hz, luminance sensitivity is typically relatively stronger, although chromatic contributions may still be present (Gelfand & Horwitz, 2018; Dobkins, Lia, & Teller, 1997). Using both frequencies therefore provides complementary leverage on possible mechanism-specific circadian effects while avoiding an all-or-none interpretation of channel isolation.

### Statistical analysis

To investigate circadian rhythms in psychophysical test variables, we employed a comprehensive multi-step statistical approach involving individual modelling of salivary melatonin concentration as a standard circadian phase marker (Duffy et al., 2011; Wang et al., 2022), and hierarchical inference on the outcome variables, temporal contrast sensitivity values of psychophysical tests directed at different retinal mechanisms. First, we fitted a circadian model to the salivary melatonin data, using a sine-cosine parameterisation with a fundamental frequency and second harmonic frequency. This allowed us to identify the best-fitting near-24 hour frequency for each participant based on the salivary melatonin, thereby approximating a person-specific endogenous period of the circadian rhythm. Next, we used this person-specific *τ* to model the psychophysical data, which covered 40 hours. We set up a Bayesian hierarchical model with sine and cosine components for the fundamental frequency, calculated using standardised clock time and the person-specific *τ*, as regressors. This model was then fitted to the psychophysical data, assuming that the data were clustered by participants. The psychophysical data included contrast sensitivity measurements from different tests probing temporal contrast sensitivity to different silent-substitution stimuli.

#### Preprocessing data

We transformed the stimulus contrast thresholds obtained from the staircase procedure into contrast sensitivity values by taking the reciprocal. To facilitate group analysis, timestamps of individual assessments were adjusted to a common reference time (07:00 as wake-up time and 08:00 as experimental start), thereby aligning the test schedule of all participants regardless of their actual wake-up times. The complete dataset did not contain any missing values. In case of device failures that disrupted data collection, we immediately repeated the respective test to ensure the integrity and completeness of the dataset.

#### Model selection

The foundation of all statistical models is the assumption of a cosinor model, defined as:

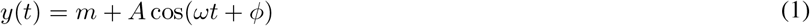

where *y* is the dependent outcome variable, *m* is the mean average intercept or MESOR (Midline Estimating Statistic Of Rhythm), *A* is the amplitude, *ω* is the angular frequency corresponding to a circadian period that is typically close to *τ* = 24 hours, *t* is time, and *ϕ* is the phase shift. Based on the trigonometric angle sum identity, the model can be rewritten as a sine-cosine parameterisation:

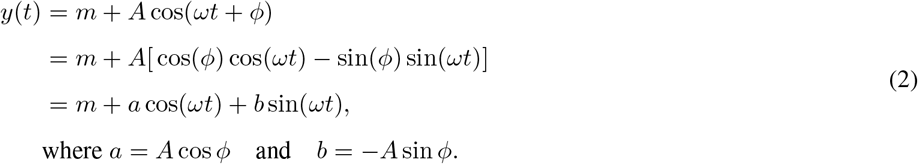

This parameterisation is preferred because it expresses oscillatory components directly instead of amplitude and phase, making it easier to estimate parameters, and it can be extended by sine and cosine terms to account for other effects, such as asymmetrical peaks and troughs. From a symmetrical model with fundamental frequency only, amplitude (*A*) and phase (*ϕ*) estimates of the proposed underlying circadian rhythm can be derived from the parameter estimates of the oscillatory components:

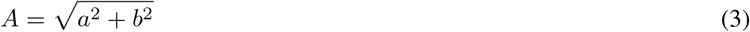

with *ϕ* ∈ (−*π, π*] corresponding to 0 to *τ* hours, *ϕ* is calculated with:

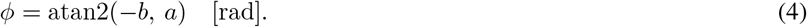

Note that due to the definition of *b* in the parameterisation in Equation 2, a negative sign is applied to *b* here. When modelling data using not only a fundamental frequency but also one or more harmonics, the amplitude and phase of each harmonic frequency are estimated separately. In hierarchical models, amplitude and phase of the group effect can be calculated as detailed above in Equations 3 and 4. To derive parameter estimates for a person-specific cosinor model, the average effect must be added to the person-specific random effect before calculating the amplitude and phase based on the sum, as these operations are non-commutative.

#### Model setup of salivary melatonin data

The aim of modelling the salivary melatonin data was to derive a participant-specific estimate for the current period of the circadian pacemaker during the observation window. Theoretical caveats to this idea are detailed in Appendix A. Therefore, model estimation was performed separately for each participant using linear regression. We employed a cosinor model in a sine–cosine parameterisation to characterise the periodic secretion of melatonin, with terms at the fundamental frequency (1/*τ*) and its second harmonic (2/*τ*). We first computed oscillatory sine and cosine regressors

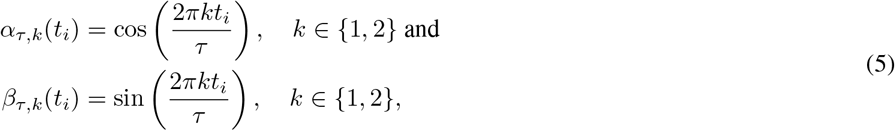

that served as periodic terms in the linear model. The statistical model for a participant’s salivary melatonin data 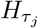 was defined as a harmonic series of *k* = 2 terms:

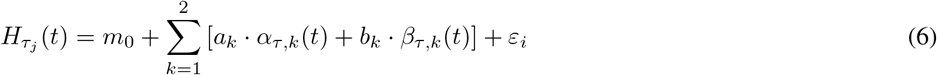

where *t* denotes hours passed since wake-up. *m*_0_ represents the average melatonin concentration in the saliva. The parameters *a*_*k*_ and *b*_*k*_ were estimated separately for each participant *j*, allowing for individual differences in amplitude contributions of both harmonics.

Finally, to estimate the participant-specific period, we conducted a grid search over *τ*_*j*_ ∈ [20, 28] with 1-minute resolution (Δ*τ* = 1/60 h). The best-fitting *τ*_*j*_ for participant *j* was selected by minimising the Akaike Information Criterion (AIC) across the grid. The linear regression models were fitted with ordinary least squares method using R’s `lm` function.

#### Model setup for psychophysical data

To fit the psychophysical contrast threshold data, we selected a linear mixed model. First, the two-level structure accounts for the clustering of data within participants, as every participant contributed around 33 single data points for temporal contrast sensitivity for each retinal mechanism. Second, linearity between a sinusoidal predictor and a sinusoidal contrast sensitivity was assumed. We used a Bayesian approach with a normal prior distribution for all model parameters. The full model assumes a data-generating process with a circular process of a certain, participant-specific length *τ*_*j*_. The full model is as follows:

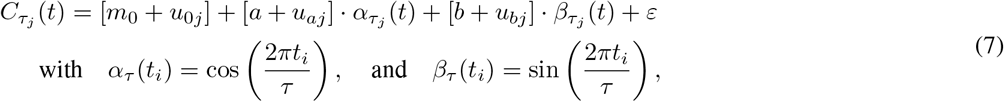

where 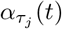 and 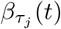 are calculated based on the participant-specific optimal *τ*_*i*_ obtained with the model fit based on Equations 6. The null model is a fixed intercept and random intercept model, missing the proposed circadian effect, thus explaining the data with only participant-specific constants. The null model was specified as:

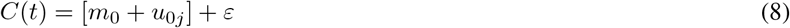

where *m*_0_ is the average intercept (or MESOR), *u*_0*j*_ is the participant-specific deviation of this intercept; *α*_*j,i*_ and *β*_*j,i*_ together indicate the shape amplitude of a circadian rhythm that varies over time *t*. Further, the model assumes that the data are clustered within a participant with participant-specific random effects: *u*_0*j*_ is the participant-specific random intercept, *u*_*aj*_ and *u*_*bj*_ are the residuals of the random slope for each participant *j* without the fixed effect. Finally, *ε* is the error term at each response *i* at time *t* and including clustering due to multiple responses *i* coming from one participant *j*. Model estimations are carried out with the `brm` function of the `brms` package (Bürkner, 2017).

#### Statistical inference

To evaluate the hypotheses, we compared full models to null models using the Bayes Factor (*BF*). The Bayes Factor is a metric used to quantify the strength of evidence in favour of one statistical model over another. The null model, as defined in Equation 8, assumes that temporal contrast sensitivity remains stable over time for each individual. It incorporates a fixed intercept and a random intercept to account for differences in baseline temporal contrast sensitivity between individuals. The full model, as described in Equation 7, assumes that temporal contrast sensitivity not only differs between individuals but also co-varies with the individual’s circadian rhythm (a fixed effect). Additionally, the full model allows for variation in this relationship across participants by including random slopes, alongside random intercepts, to capture differences in how individuals’ temporal contrast sensitivity fluctuates over time.

The Jeffreys scale is used as an interpretation guideline for the level of support for different ranges of BF as listed in Table 4 (Jeffreys, 1998). Here, the BF quantifies the strength of evidence for the full model, implying an underlying circular process with a period of *τ* . Categorisation of the Bayes factor and the natural logarithm of the Bayes factor (ln BF) is interchangeable, e.g. BF > 100 corresponding to ln BF > 4.61.

**Table 4:** Interpretation of Bayes Factor values and their corresponding natural logarithm values.

| Bayes Factor (BF) | Interpretation | ln of BF |
| --- | --- | --- |
| $BF > 100$ | Decisive evidence for the full model | $\ln BF > 4.61$ |
| $30 < BF < 100$ | Very strong evidence for the full model | $3.40 < \ln BF < 4.61$ |
| $10 < BF < 30$ | Strong evidence for the full model | $2.30 < \ln BF < 3.40$ |
| $3 < BF < 10$ | Moderate evidence for the full model | $1.10 < \ln BF < 2.30$ |
| $1 < BF < 3$ | Weak evidence for the full model | $0 < \ln BF < 1.10$ |
| $\frac{1}{3} < BF < 1$ | Weak evidence for the null model | $-1.10 < \ln BF < 0$ |
| $\frac{1}{10} < BF < \frac{1}{3}$ | Moderate evidence for the null model | $-2.30 < \ln BF < -1.10$ |
| $\frac{1}{30} < BF < \frac{1}{10}$ | Strong evidence for the null model | $-3.40 < \ln BF < -2.30$ |
| $\frac{1}{100} < BF < \frac{1}{30}$ | Very strong evidence for the null model | $-4.61 < \ln BF < -3.40$ |
| $BF < \frac{1}{100}$ | Decisive evidence for the null model | $\ln BF < -4.61$ |

For lack of prior evidence, we abstain from adding further covariates, such as age and sex in the confirmatory analysis. We use posterior sampling to obtain the posterior distribution of the model parameters. Inference about the presence of a circadian modulation of temporal contrast sensitivity is drawn based on conventions in interpreting the Bayes Factor obtained during model comparison, and we use the `posterior_summary()` function from the `brms` package to sample the posterior distribution of the model parameters. We sample from the posterior distributions to obtain parameter estimates to derive amplitude and phase.

#### Statistical implementation

While some preprocessing steps were implemented in Python version 3.10, statistical models were estimated in R version 4.4.3. All data and analysis code used in this manuscript are openly available in a GitHub repository, released under the MIT licence, at https://github.com/tscnlab/HeinrichsEtAl_JVis_2025.

## Results

In the following, we report summary and descriptive statistics for the variables used, e.g. salivary hormone concentration and psychophysical threshold, as well as inferential statistics for our confirmatory hypothesis. For all model comparisons, the obtained BF indicates the strength of evidence for the full model, which included the circular components, when compared with the null model, which assumes no circular process. For better readability, we report the natural logarithm of the Bayes factor (ln BF) instead of the BF in the results section.

### Descriptive statistics

#### Salivary melatonin concentrations

Salivary melatonin concentrations ranged from 0.0 to 46.8 picograms per milliliter (pg/ml). The overall mean melatonin concentration was 7.06 ± 9.05 pg/ml, whereas the median was 2.64 pg/ml. The difference between mean and median suggests a non-normal distribution of overall melatonin levels. Indeed, the distribution of concentration levels across time and all participants for salivary melatonin is a zero-inflated distribution, see Figure 2.

##### Individual and group dynamics in melatonin concentration

Onset of melatonin secretion as well as concentration peaks vary between participants, as can be seen in the participant-specific plots in Figure 5, panels C and D (see below). The average melatonin profile plotted in Figure 4B reflects time-restricted melatonin secretion: across the sample, there are consistently low or null concentration values during habitual daytime, with small standard deviation. This is also shown by the strong accumulation of near-zero samples in Figure 2A. For 6 participants, at some point during the day, salivary melatonin concentration did not exceed the detection threshold, as indicated by zero values; for the other 6 participants, the individual minimum salivary melatonin levels ranged from 0.1 to 2.2 pg/ml. There is a broad peak in concentration levels starting a few hours before habitual bedtime, extending across the night, and dropping before habitual wake time. Daytime levels are re-established shortly after habitual wake time. The dynamics of individual salivary melatonin levels throughout the study protocol are visually consistent with the average melatonin profile: For all participants, melatonin concentrations initially decrease as the protocol starts one hour after habitual wake-up. Thereafter, the participant-specific melatonin profiles all exhibit low levels during subjective daytime, dropping close to zero with an average of 1.26 pg/ml by the start of the second cycle (approx. 3.5 hours after wake-up) and an average daytime value (defined as time between habitual wake-up and habitual bedtime) of 3.69 pg/ml. A similar drop can be seen by the seventh experimental block (mean concentration of 1.53 pg/ml) which again corresponds to 3.5 hours after habitual wake time. At subjective nighttime, average melatonin levels rise to an average of 17.76 ± 9.17 pg/ml for the time between habitual bedtime and habitual wake-up. Note that habitual daytime and habitual nighttime refer to pre-session stabilisation and do not reflect the sleep–wake or rest–activity behaviour during the experiment, which took place on a 3.75-hour rhythm (corresponding to the experimental block). The course of concentration levels for the second night is cut off 2 hours after habitual bedtime. Peak concentration levels do not seem to differ greatly between the first and second habitual nights; the average concentration at the beginning of the first night is 20.57 ± 9.86 pg/ml while the average concentration at the beginning of the second night is 18.81 ± 9.18 pg/ml.

**Figure 4.**
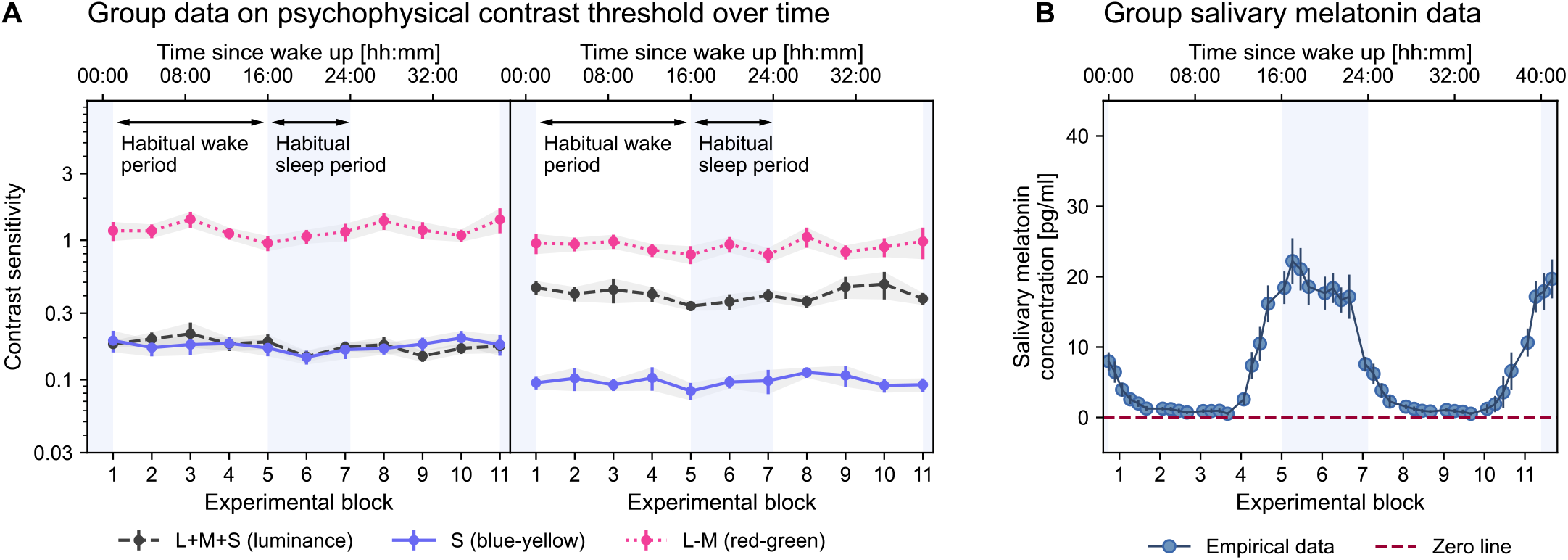
Participant-averaged time courses of temporal contrast sensitivity and salivary melatonin concentration. Temporal contrast sensitivity is shown separately for the 2 Hz and 8 Hz conditions, with different curves for the three stimulus directions. Curves represent participant means and ribbons denote the standard error of those participant means. Vertical shaded bands indicate habitual sleep times. Data are plotted in relative clock time, with wake time used as the alignment reference.

**Figure 5.**
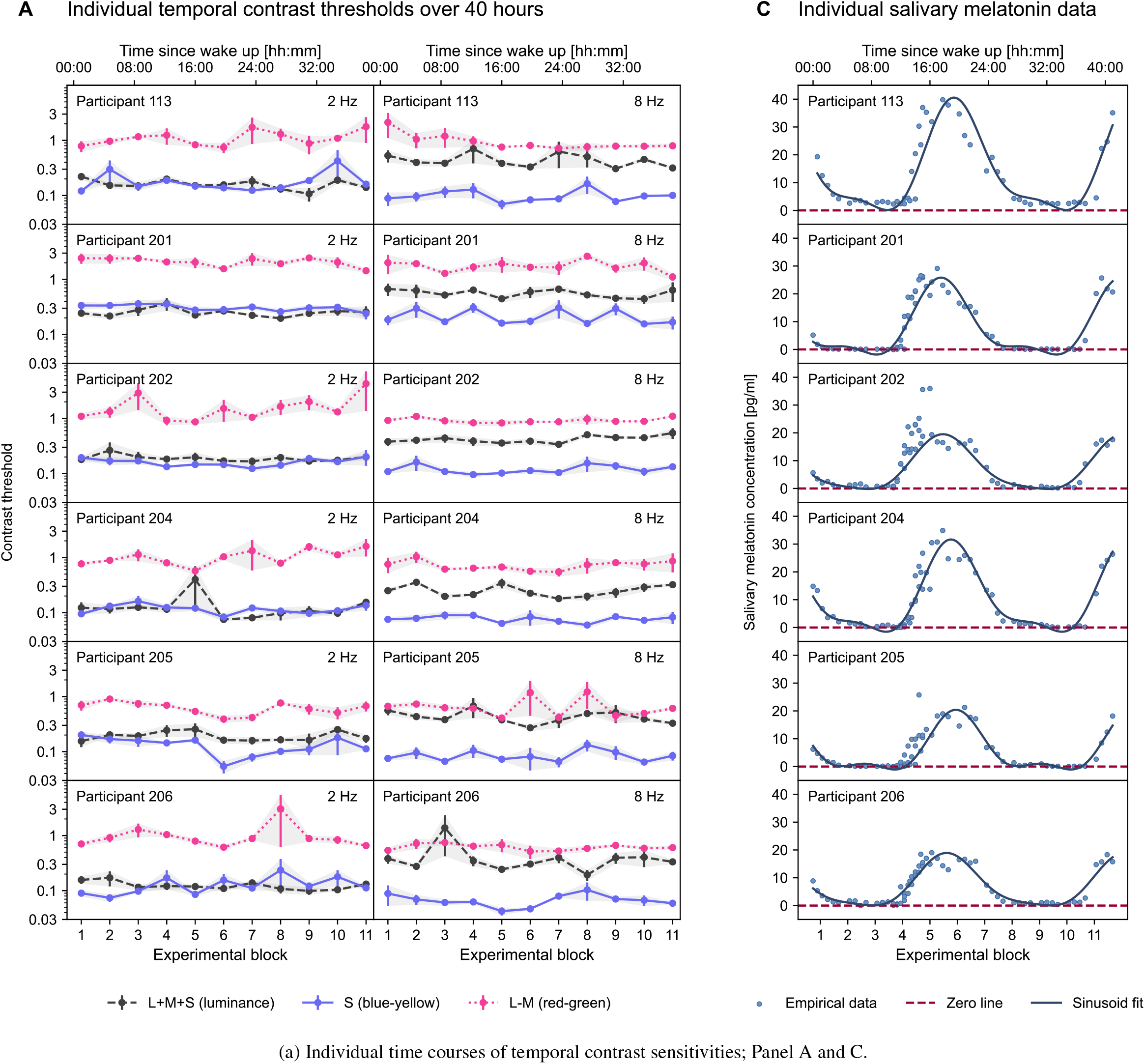
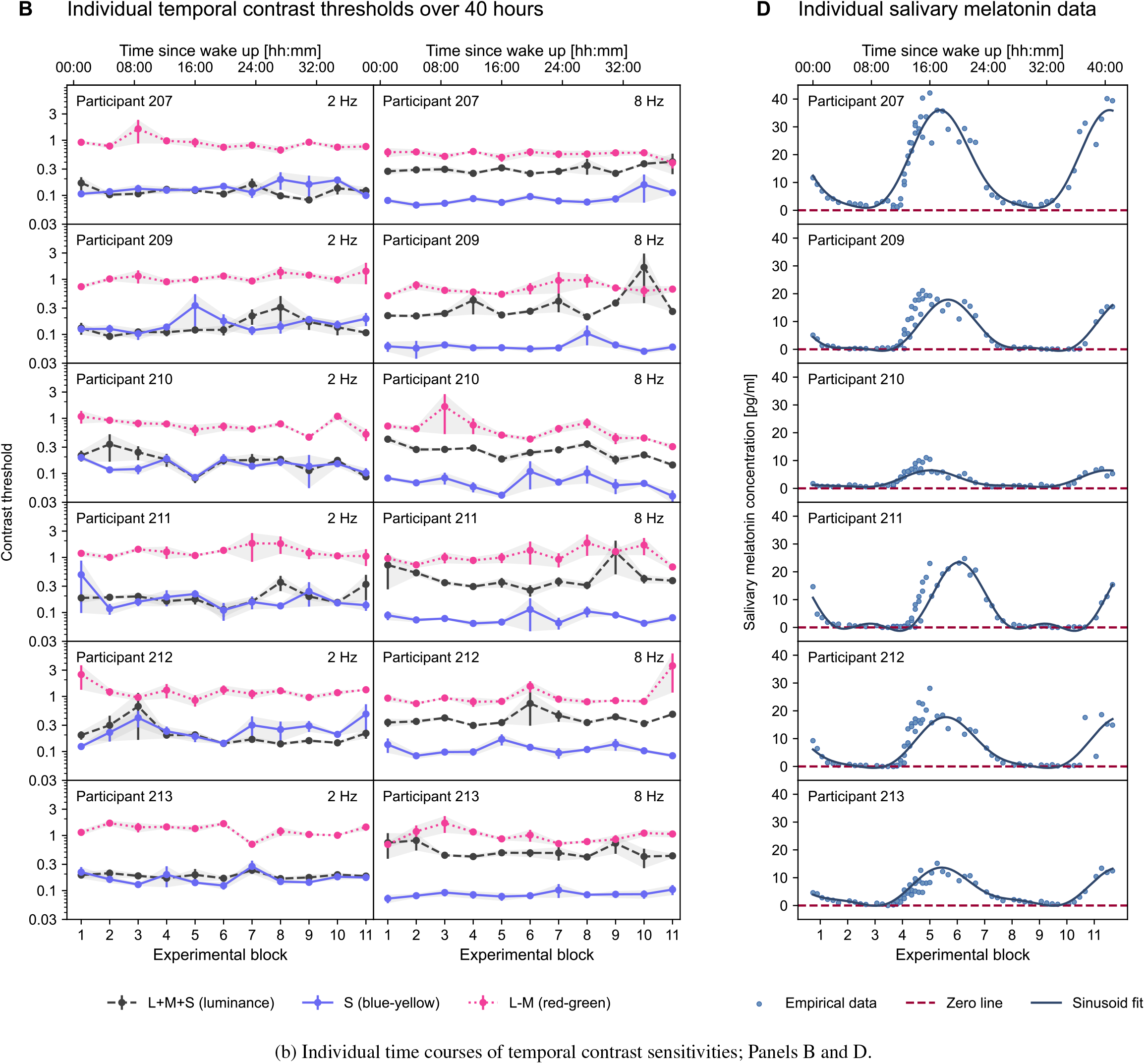
Individual time courses of temporal contrast sensitivities; panels A and C (previous page) and panels B and D (current page). Individual time courses of temporal contrast sensitivities; one row per participant, two columns for two different temporal frequencies. (A and B) Psychophysical contrast sensitivity estimates of complete study sample split into two subfigures. Each point marks the mean contrast sensitivity across all tests within one experimental block, shading indicates standard deviation of the mean. (C and D) Melatonin concentration of the complete study sample split into two subfigures, including the fit of a near-24-hour rhythm; participant-specific *τ* was derived from the linear model showing the maximum *AIC*.

##### Inter-individual variability in melatonin profiles

Participants differed in their average concentration (7.07 ± 3.35 pg/ml) likely reflecting differences in the range of melatonin secretion, peak levels, or duration of melatonin secretion (i.e. elevated salivary melatonin levels). All participants exhibited time-dependent variation in their melatonin profiles with discernible peaks. The highest peak occurred in participant 113 (46.8 pg/ml) and was sixfold higher than the lowest peak observed in participant 210(7.55 pg/ml). Accordingly, the within-subject range (maximum - minimum) also varied across participants (mean 24.16 ± 10.72 pg/ml).

#### First-stage model fitting of melatonin data

There is variability between participants in maximum levels and overall volume of salivary melatonin and, to some extent, also in the timing of salivary melatonin secretion across the day, suggesting that these dynamics can be described by a circular process with participant-specific amplitude and phase. Due to the zero-inflated distribution of the concentration values, we refitted a multi-harmonic model with the different fundamental frequencies and the respective second harmonic as described in section Statistical analysis.

##### Melatonin derived circadian period

Optimal fundamental frequency 1/*τ* was determined by a grid search across different fundamental frequencies, maximising model fit as indicated by the Akaike Information Criterion (*AIC*). The mean *AIC* for all identified participant-specific models is 206.19. For visualisation, Figure A2 in Appendix A plots the coefficient of determination (R^2^) as a function of *τ*, with the optimal participant-specific *τ* indicated. The resulting model fits for each individual are visualised above the raw data in Figure A1 in Appendix A. For example, the salivary melatonin concentration of participant 201 is best described by the sum of signals at 1/24.63 = 1.13 × 10^−5^ Hz and 2/24.63 = 2.26 × 10^−5^ Hz. The mean optimal *τ* of the sample is 24.18 ± 0.39 hours, revealing a near 24-hour rhythm for all participants. The participant-specific *τ* values range from 23.22 hours as the shortest period for participant 207 to 24.65 hours as the longest period for participant 202.

##### Oscillatory parameters of melatonin rhythm

Mesor, as well as the amplitudes and phases of the fundamental frequency and second harmonic describing the participant-wise melatonin concentration, are listed in Table 5. From the final model, both amplitude and phase of the first (fundamental) and second harmonic were derived from the first and second set of sine-cosine parameter sets, respectively. Circular statistics derived from the fundamental components are visualised in the polar plot in Figure 2B. The amplitudes of the fundamental melatonin rhythms measure on average 10.87 ± 4.55 pg/ml, ranging from 2.81 to 18.8 pg/ml. Average phase for the fundamental is 1.46 ± 0.28 (05:34±01:05 hours), ranging from 1 (03:50 hours) to 2.03 (07:46). Conversion into hours of phase angle of the fundamental component are based on an ideal 24:00 hour sinusoidal rhythm. Amplitudes of the second harmonic were at 3.45 ± 1.62 pg/ml, ranging from 0.94 to 6.12 pg/ml. The distribution of the different phases is quite narrow. The average phase of the second harmonic is 4.87 ± 2.77, ranging from 2.84 to 3.04 radians. With a lower peak amplitude than the fundamental, the second harmonic may not drastically change the dynamics of melatonin concentration, but may slightly change the timing of the peak of the melatonin rhythm, as can be seen in Figure 2C.

**Table 5:** Results from best fitting participant-specific model on saliva melatonin data when iterating over *τ* s; optimal *τ* and *AIC*, as well as calculated circadian parameters listed.

| Participant | Optimal tau | $AIC$ | Intercept | Fundamental | | 2nd harmonic | |
| --- | --- | --- | --- | --- | --- | --- | --- |
|  |  |  |  | Amplitude (1st) | Phase (1st) | Amplitude (2nd) | Phase (2nd) |
| 113 | 24.48 | 284.80 | 15.83 | 18.80 | 1.23 | 6.12 | 2.84 |
| 201 | 24.63 | 235.21 | 8.55 | 12.61 | 1.72 | 4.76 | -2.52 |
| 202 | 24.65 | 201.16 | 7.68 | 9.63 | 1.68 | 2.26 | -2.62 |
| 204 | 24.07 | 234.65 | 11.31 | 15.31 | 1.25 | 5.20 | 2.88 |
| 205 | 24.10 | 178.99 | 6.52 | 9.73 | 1.14 | 4.17 | 2.43 |
| 206 | 24.35 | 211.76 | 7.58 | 9.22 | 1.49 | 2.18 | -2.99 |
| 207 | 23.22 | 266.96 | 15.13 | 17.36 | 1.52 | 3.62 | -2.98 |
| 209 | 24.22 | 183.41 | 6.17 | 8.75 | 1.43 | 2.98 | 3.04 |
| 210 | 24.28 | 126.68 | 2.77 | 2.81 | 2.03 | 0.94 | -1.92 |
| 211 | 23.92 | 164.91 | 7.25 | 11.06 | 1.00 | 5.15 | 2.08 |
| 212 | 23.90 | 233.47 | 6.84 | 8.89 | 1.42 | 2.07 | -3.10 |
| 213 | 24.32 | 152.24 | 5.53 | 6.30 | 1.57 | 1.96 | -2.55 |
| $M(SD)$ | 24.18 (0.39) | 206.19 (47.21) | 8.43 (3.84) | 10.87 (4.55) | 1.46 (0.28) | 3.45 (1.62) | -0.45 (2.77) |

#### Temporal contrast sensitivity

Temporal contrast thresholds were determined adaptively with staircase procedures. Each test targeted one stimulus direction and contained two interleaved staircases, one for 2 Hz and one for 8 Hz temporal modulation. Originally, three repetitions per stimulus direction and wake period were planned, corresponding to approximately 33 tests per direction and participant across the 11 wake periods. Occasional software failures during data transfer from the control tablet to the acquisition computer required some tests to be repeated. Conversely, a small number of superfluous repetitions or omissions arose from human error in test selection or repetition handling (cf. Table 6). Across *N* = 12 participants, 1181 tests were completed, corresponding to 2362 frequency-specific staircase estimates. Participants contributed between 31 and 34 tests per stimulus direction across the protocol, and between 93 and 100 tests in total.

**Table 6:** Deviations in test count for temporal contrast sensitivity in each colour direction. Each participant needed to complete three tests per colour direction per block. Table only contains test count per participant per experimental blocks, when there were either more or less than three tests in at least one colour direction.

| $N_{\text{Tests}}$ | Participant ID | Experimental Block | Postreceptoral mechanism | | |
| --- | --- | --- | --- | --- | --- |
|  |  |  | L+M+S (luminance) | S (blue-yellow) | L–M (red-green) |
| 6 | 113 | 0 | 2 | 2 | 2 |
| 7 | 113 | 8 | 2 | 3 | 2 |
| 8 | 113 | 9 | 3 | 2 | 3 |
|  | 202 | 1 | 3 | 3 | 2 |
|  | 206 | 4 | 3 | 2 | 3 |
|  | 211 | 1 | 3 | 3 | 2 |
|  | 211 | 9 | 2 | 3 | 3 |
|  | 212 | 1 | 3 | 3 | 2 |
|  | 212 | 9 | 3 | 3 | 2 |
|  | 213 | 6 | 3 | 3 | 2 |
|  | 213 | 10 | 3 | 3 | 2 |
| 10 | 201 | 1 | 3 | 4 | 3 |
|  | 202 | 6 | 3 | 4 | 3 |
|  | 202 | 8 | 3 | 3 | 4 |
|  | 204 | 7 | 3 | 3 | 4 |
|  | 206 | 1 | 4 | 3 | 3 |
|  | 209 | 5 | 3 | 4 | 3 |
|  | 212 | 2 | 3 | 3 | 4 |

##### Example staircase trajectories

Figure 3 shows representative staircase data from one participant and one experimental block. The figure is intended to illustrate the adaptive procedure rather than to establish any inferential pattern. For each trial, the plotted value is the presented stimulus contrast, and symbol fill indicates whether the response was correct or incorrect. Threshold estimates were obtained only after staircase termination, from the reversal values described in the Methods section. The example traces illustrate the expected qualitative range of thresholds across stimulus directions. Thresholds for the L–M condition were lowest in the displayed example, whereas the L+M+S and S-cone-directed conditions required higher nominal contrast. As in the full dataset, the relative ordering differed across stimulus directions and temporal frequencies, underscoring that the figure is descriptive only.

##### Group averages in contrast sensitivity

Contrast sensitivity was calculated as the reciprocal of stimulus contrast threshold. Table 7 summarises the grand means across participants and time. Because the number of tests per block was not perfectly balanced, we first averaged repeated tests within participant and block and then computed group means from those participant-level block means so that each participant contributed equally. Across the dataset, contrast sensitivity was highest for the L–M condition and lower for the L+M+S and S-cone-directed conditions. The relative difference between 2 Hz and 8 Hz also depended on stimulus direction, consistent with the known temporal tuning differences between chromatic and luminance-sensitive mechanisms. These descriptive differences motivated the use of condition-specific inferential models rather than a single pooled analysis across all stimulus directions and frequencies.

**Table 7:** Descriptive statistics of temporal contrast sensitivity tasks; average contrast sensitivities per colour condition and frequency across participants and time.

| Postreceptoral channel | Temporal frequency [Hz] | Temporal contrast sensitivity |  |
| --- | --- | --- | --- |
|  |  | Mean | Standard deviation |
| S (blue-yellow) | 8 | 0.098 | 0.062 |
|  | 2 | 0.175 | 0.118 |
| L+M+S (luminance) | 8 | 0.410 | 0.341 |
|  | 2 | 0.177 | 0.136 |
| L–M (red-green) | 8 | 0.903 | 0.654 |
|  | 2 | 1.191 | 0.870 |

##### Group-level time course in temporal contrast sensitivity

If temporal contrast sensitivity were modulated by circadian phase, one would expect a systematic near-24-hour fluctuation in the aligned group time courses. Visual inspection of Figure 4 does not reveal such a pattern. Variation in block-wise group means was small, and any apparent drifts were minor relative to the uncertainty around the means. It therefore remains possible that small participant-specific dynamics are present, but the group averages do not show an obvious circadian signature.

##### Participant-level time course in temporal contrast sensitivity

Participant-level time courses broadly resembled the group summaries. Figure 5 shows variability across individuals, stimulus directions, and frequencies, but no consistent pattern that would support a common circadian phase relationship across participants. Some participants exhibited relatively stable performance across blocks, whereas others showed isolated excursions in single blocks. Because only a small number of tests contributed to each block average, these excursions can influence descriptive plots disproportionately. For statistical inference, we therefore modelled the single-test data directly rather than relying on block averages.

### Inference statistics

#### Temporal contrast sensitivity data

In the confirmatory analyses, contrast sensitivity was log-transformed with the natural logarithm (ln) to better approximate a normal distribution. Derived amplitudes are therefore reported in log units.

##### Confirmatory tests on “L–M (red-green)” mechanism

Model comparisons were conducted separately for “L–M (red-green), 2 Hz” and “L–M (red-green), 8 Hz” (coefficients in Table 8, fit statistics in Table 9). Both conditions showed the same overall pattern: decisive evidence in favour of the null model and only very small oscillatory estimates. For “L–M (red-green), 2 Hz”, parameter estimates of *α* = 0.04 and *β* = −0.02 corresponded to a group-level amplitude of 0.04 log units in contrast sensitivity. This was the only model in which the 95% credible interval for one fixed effect (*α*) did not include zero, but the model comparison still decisively favoured the null model (ln BF = −11.76). The corresponding group-level phase was −0.45 rad. Accounting for participant-level variation in the sine (*SD*_*α*_ = 0.02) and cosine (*SD*_*β*_ = 0.02) components yielded individual phase estimates ranging from −2.21 to −1.75 rad. For “L–M (red-green), 8 Hz”, the oscillatory estimates were similarly small. Parameter estimates of *α* = 0.03 and *β* = −1.74 × 10^−3^ yielded a group-level amplitude of 0.03 log units and a phase of −0.06 rad. The 95% credible intervals for both fixed effects included zero, and the model comparison again decisively favoured the null model (ln BF = −11.76). Conditional *R*^2^ values differed only minimally between the full model 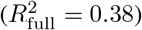 and the null model 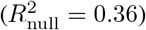, indicating that the oscillatory terms explained little additional variance.

**Table 8:**
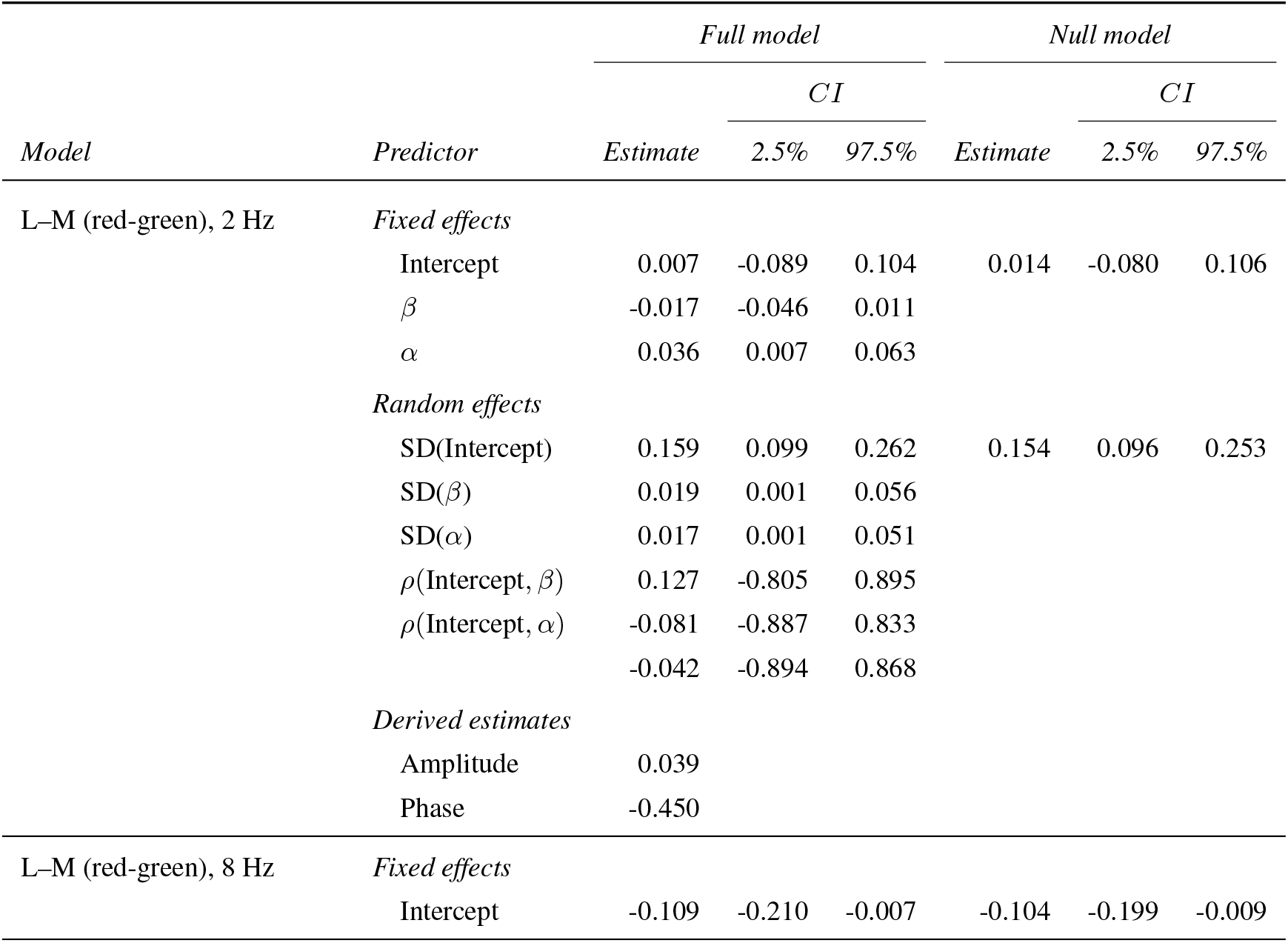
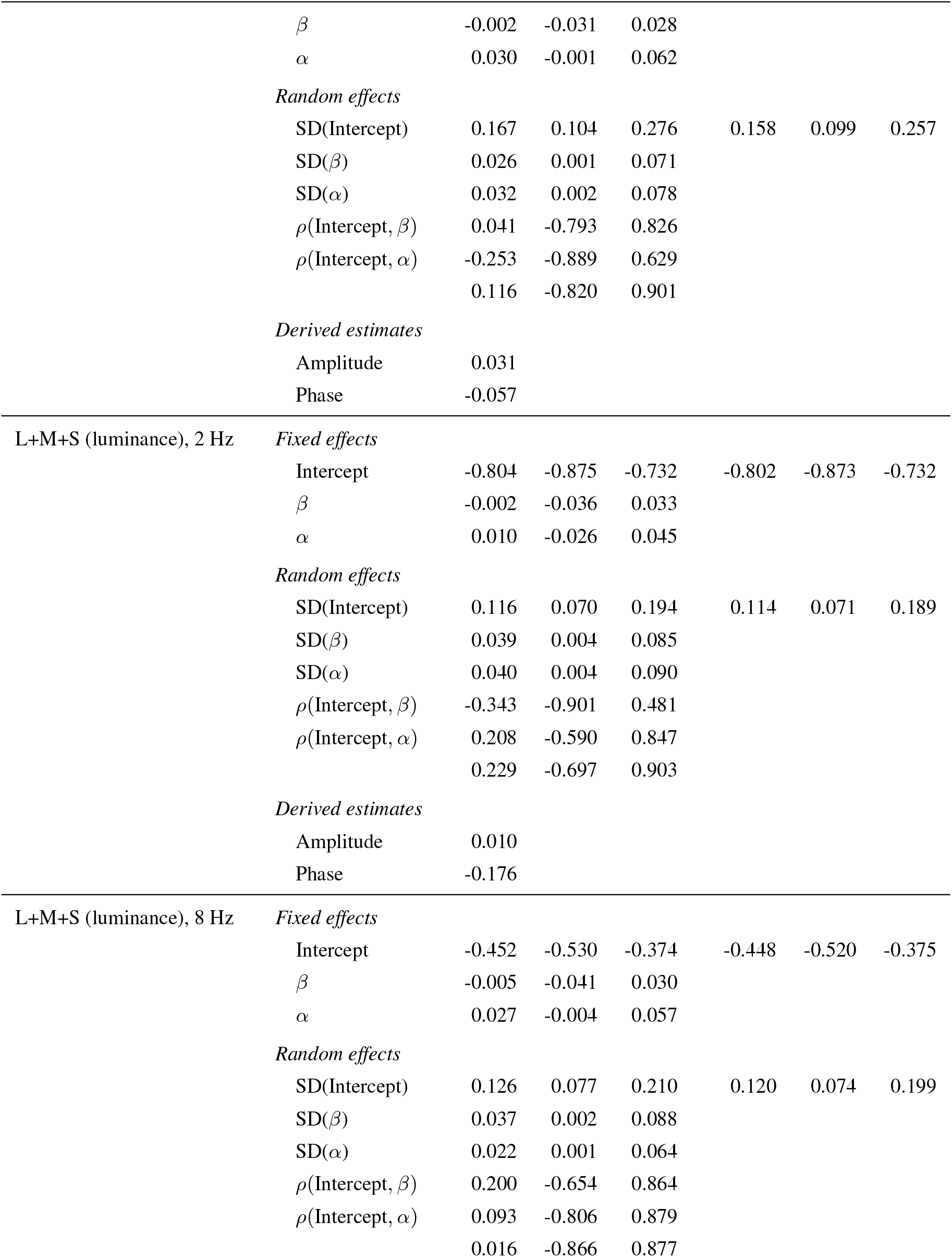
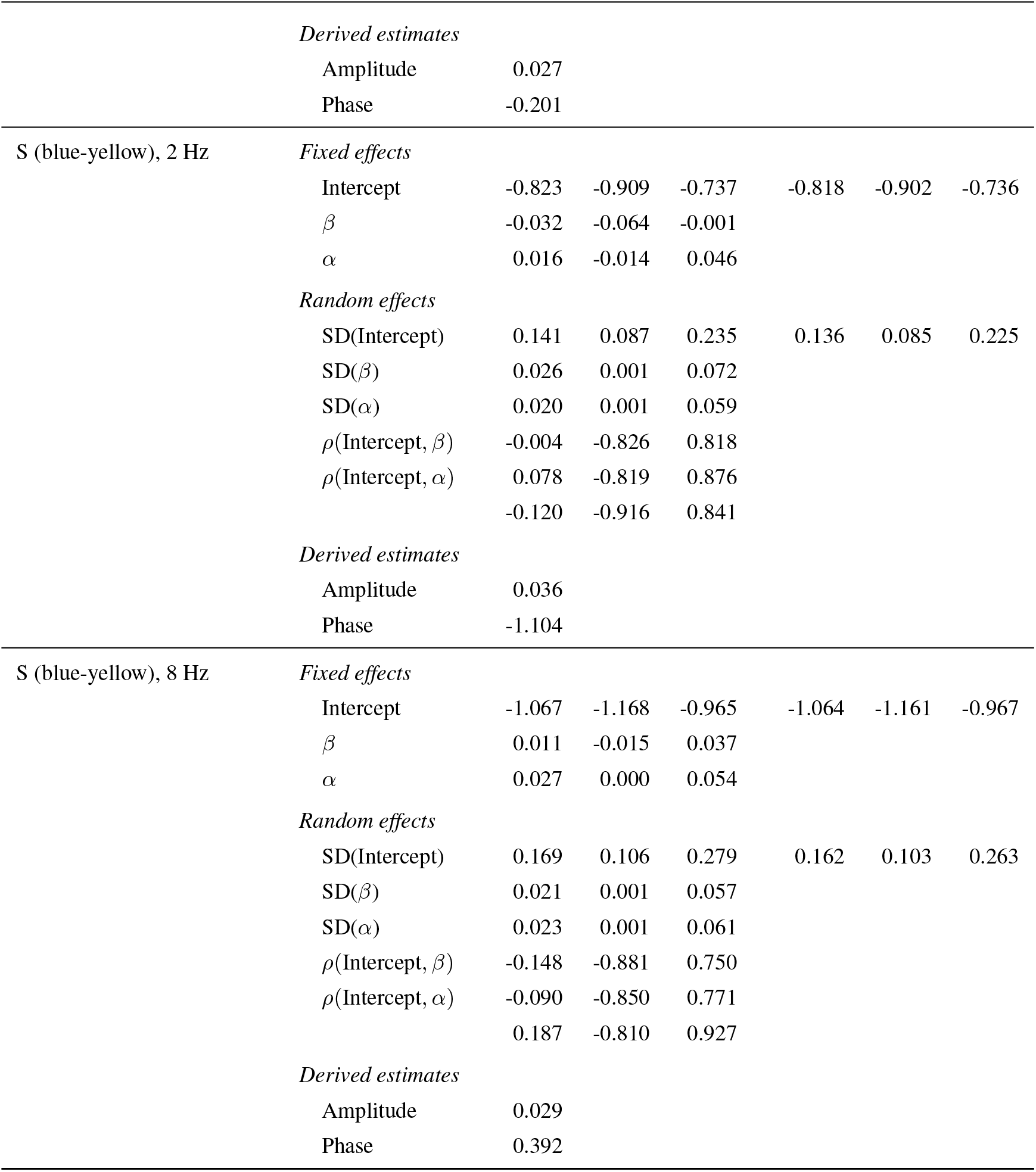
Fixed- and random-effect estimates and derived group-level amplitude and phase for all six condition-by-frequency models. Within each model block, the left *Estimate*/*CI* columns refer to the full (circadian) model and the right columns to the null model. Fit statistics are reported separately in Table 9.

**Table 9:** Sample sizes, model-comparison evidence and explained variance for all six models. Conditional and marginal *R*^2^ are reported for the full (circadian) and null models. Coefficient estimates are reported in Table 8.

| *Model* | $N_{\text{Obs}}$ | $N_{\text{Clust}}$ | $\ln \text{BF}$ | Conditional $R^2$ | | Marginal $R^2$ | |
| --- | --- | --- | --- | --- | --- | --- | --- |
|  |  |  |  | *Full* | *Null* | *Full* | *Null* |
| L–M (red-green), 2 Hz | 394 | 12 | -12.153 | 0.376 | 0.356 | 0.018 | 0.000 |
| L–M (red-green), 8 Hz | 394 | 12 | -11.766 | 0.427 | 0.402 | 0.015 | 0.000 |
| L+M+S (luminance), 2 Hz | 391 | 12 | -13.083 | 0.292 | 0.257 | 0.009 | 0.000 |
| L+M+S (luminance), 8 Hz | 391 | 12 | -12.994 | 0.275 | 0.248 | 0.014 | 0.000 |
| S (blue-yellow), 2 Hz | 396 | 12 | -12.470 | 0.312 | 0.289 | 0.017 | 0.000 |
| S (blue-yellow), 8 Hz | 396 | 12 | -12.439 | 0.473 | 0.453 | 0.014 | 0.000 |

##### Confirmatory tests on “S (blue-yellow)” mechanism

For the S-cone-directed condition, both frequency-specific model comparisons also favoured the null model. The ln BF values were −12.46 for “S (blue-yellow), 2 Hz” and −12.43 for “S (blue-yellow), 8 Hz”, again indicating decisive evidence against the oscillatory model. For “S (blue-yellow), 2 Hz”, parameter estimates of *α* = 0.02 and *β* = −0.03 yielded a group-level amplitude of 0.04 log units and a phase of −1.1 rad. For “S (blue-yellow), 8 Hz”, parameter estimates of *α* = 0.03 and *β* = 0.01 yielded a group-level amplitude of 0.03 log units and a phase of 0.39 rad. In both models, the credible intervals for the fixed effects included zero. Conditional *R*^2^ values again differed only marginally between the full and null models. For “S (blue-yellow), 8 Hz”,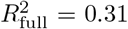 and 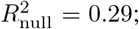 for “S (blue-yellow), 2 Hz”, 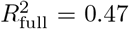 and 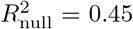. Individual amplitudes and phases varied across participants, but these differences have limited interpretability given the decisive model-comparison evidence for the null model.

##### Confirmatory tests on “L+M+S (luminance)” mechanism

We also compared the full and null models separately for “L+M+S (luminance), 2 Hz” and “L+M+S (luminance), 8 Hz” . As in the chromatic conditions, both comparisons decisively favoured the null model. The ln BF values were −13.09 for “L+M+S (luminance), 2 Hz” and −12.99 for “L+M+S (luminance), 8 Hz” . For “L+M+S (luminance), 2 Hz”, parameter estimates of *α* = 9.55 × 10^−3^ and *β* = −1.70 × 10^−3^ yielded a group-level amplitude of 9.69 × 10^−3^ log units and a phase of −0.18 rad. For “L+M+S (luminance), 8 Hz”, parameter estimates of *α* = 0.03 and *β* = −5.42 × 10^−3^ yielded a group-level amplitude of 0.03 log units and a phase of −0.2 rad. Conditional *R*^2^ values were again only slightly higher for the full than for the null models: for “L+M+S (luminance), 2 Hz”, 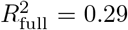 and 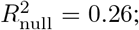 for “L+M+S (luminance), 8 Hz”, 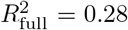 and 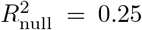. Phase dispersion was broad for “L+M+S (luminance), 2 Hz” than for the other conditions, but the estimated amplitudes remained effectively near zero therefore phase estimates may be arbitrary. Table 10 summarises the group-level amplitudes and phases as well as the corresponding individual estimates. Across “L+M+S (luminance)” and “L–M (red-green)” models, estimated amplitudes were generally in the second decimal place, and across “S (blue-yellow)” models they were typically in the third decimal place, indicating negligible oscillatory magnitudes overall.

**Table 10:**
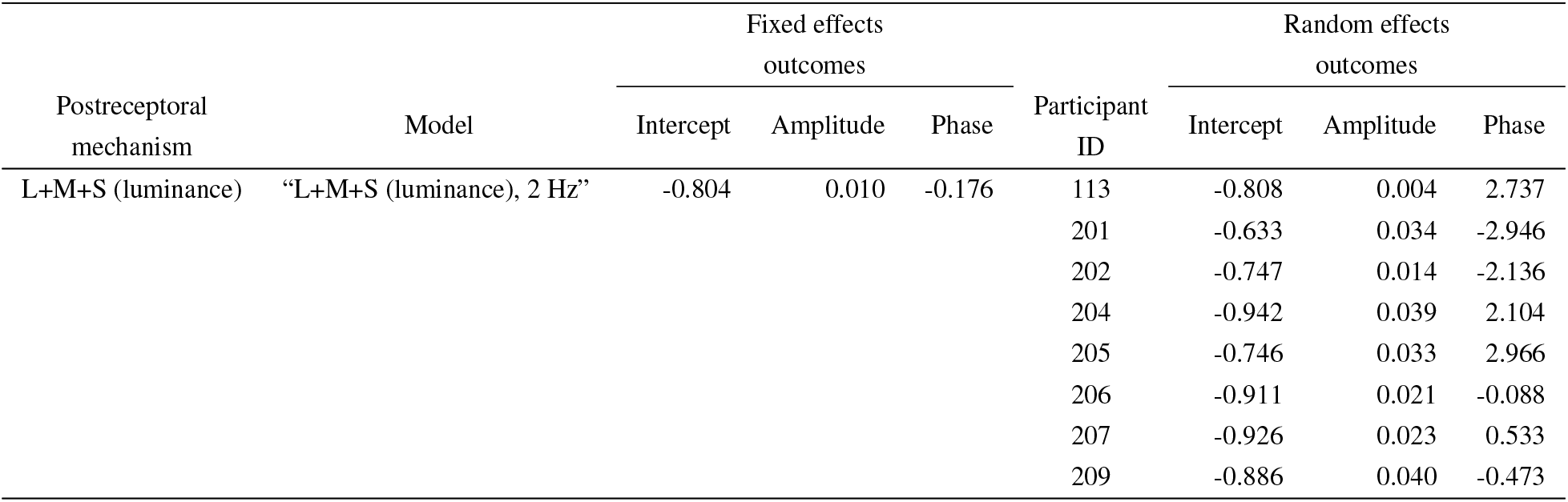
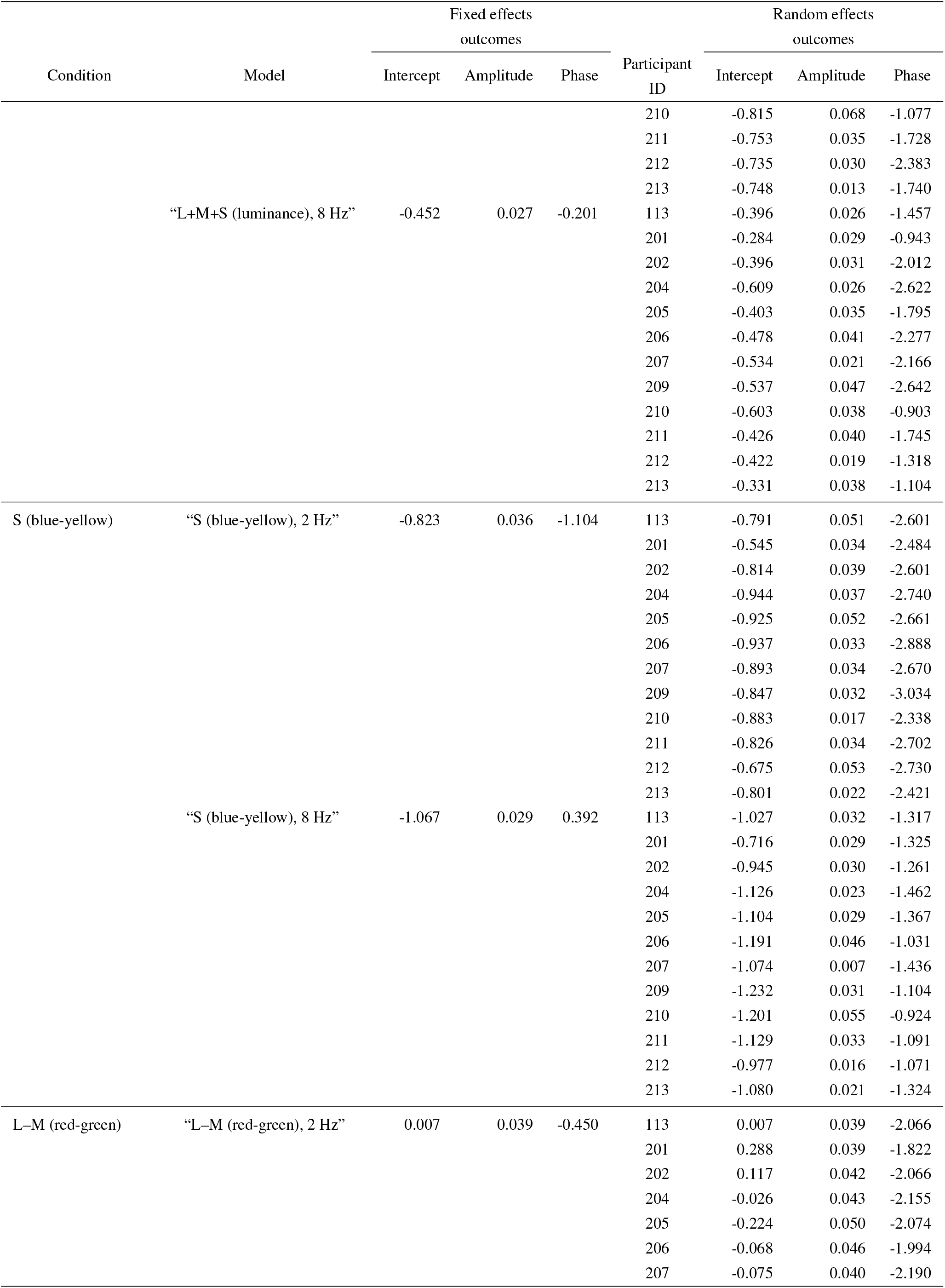
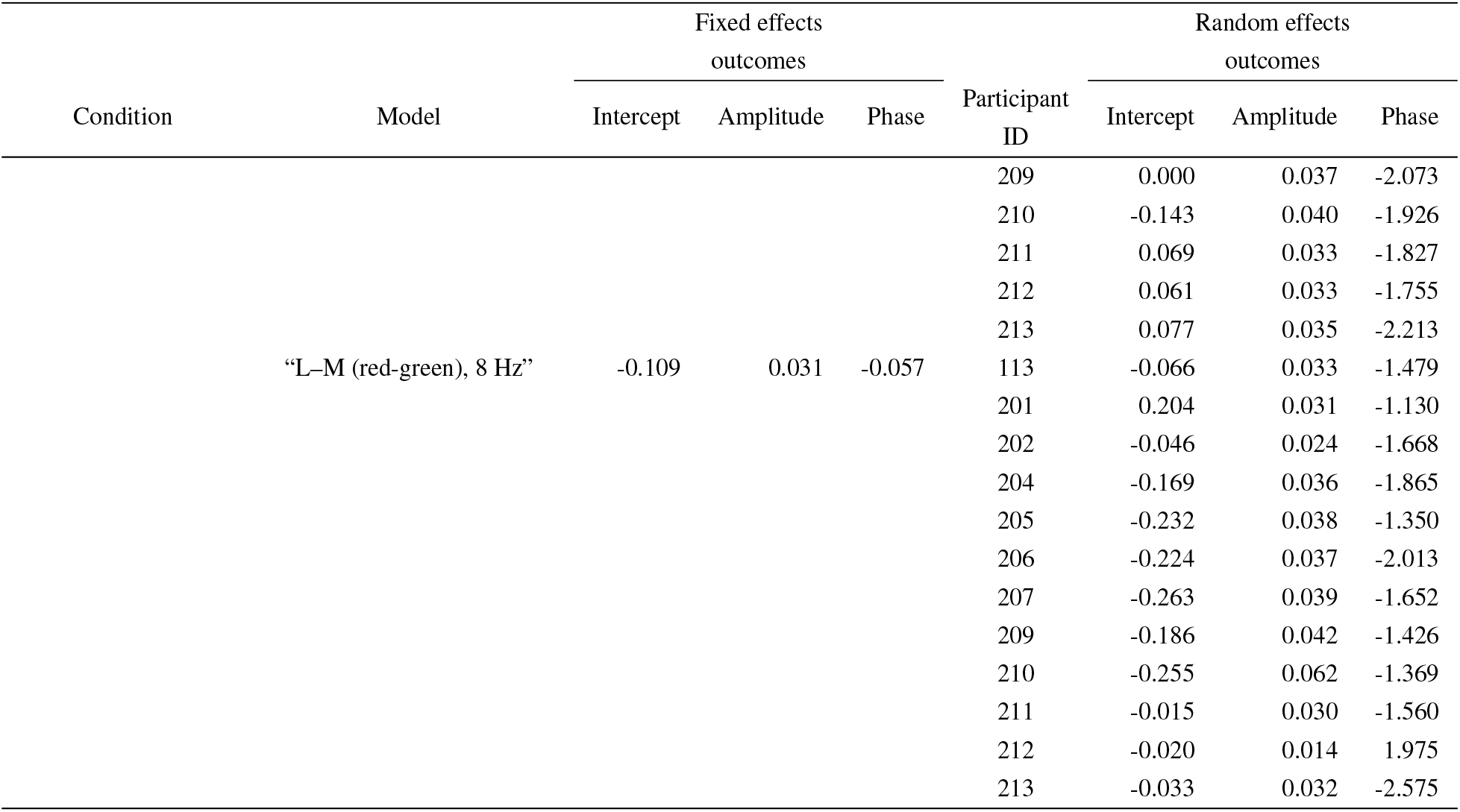
Average and participant-wise mesor, amplitude and phase based on participant-specific periods.

##### Summary

Across all six conditions, ln BF values were consistently negative and large in magnitude, ranging from −13.09 to −11.76. On Jeffreys’ scale, these values constitute decisive evidence in favour of the null model. Conditional *R*^2^ values differed only marginally between the full and null models, indicating that most explained variance reflected between-participant differences in mean contrast sensitivity rather than oscillatory variation over time.

In the full models, fixed-effect estimates for the oscillatory components were small, and their 95% credible intervals generally included zero, with one exception. The corresponding amplitudes were near zero, and phase estimates were broadly dispersed across participants. Although posterior uncertainty was not propagated into the derived amplitude and phase estimates, the overall pattern provides no evidence for a robust circadian modulation of temporal contrast sensitivity in any of the six tested conditions.

## Discussion

The present study tested whether human visual performance is modulated by the circadian pacemaker. Using a 40-hour forced desynchrony protocol with ultra-short sleep–wake cycles, we decoupled circadian phase from homeostatic state and repeatedly measured cone-mediated temporal contrast sensitivity under photopic viewing conditions. The stimuli targeted L+M+S (luminance), L–M (red-green), and S (blue-yellow) post-receptoral mechanisms at two temporal frequencies, 2 and 8 Hz. Circadian phase was indexed by salivary melatonin, and each mechanism-frequency combination was analysed with hierarchical models that either included an oscillatory component matched to the participant’s melatonin-derived period or assumed participant-specific constant levels.

Across all six mechanism-frequency combinations, model comparisons decisively favoured the constant model over the oscillatory alternative. Gains in conditional R^2^ were minimal, fixed-effect estimates were small with credible intervals largely including zero, and derived amplitudes were near zero with phases broadly dispersed across participants. Taken together, these results indicate no detectable near-24-hour modulation of cone-mediated temporal contrast sensitivity under constant ambient conditions designed to isolate circadian from homeostatic influences.

### Interpretation and context

Our null results are unexpected in light of prior reports of time-of-day differences in visual sensitivity and of circadian modulation of retinal physiology (O’Keefe & Baker, 1987; Roenneberg, Lotze, & Von Steinbüchel, 1992). Earlier studies have examined diurnal effects in a range of visual outcomes, including increment thresholds and luminance-contrast tasks (Andrade, 2022; Andrade, Neto, et al., 2018; Andrade, Cristino, et al., 2018; Roenneberg et al., 1992). A key difference between those studies and the present one lies in the inferential target: many time-of-day studies quantify visual variation under habitual sleep–wake schedules, where circadian phase strongly co-varies with time awake and other behavioural variables relevant to homeostatic regulation. Such designs cannot determine whether observed variation is circadian in origin or instead reflects homeostatic correlates such as sleep pressure, activity, posture, or hunger. The possibility of circadian influence is often discussed, but the underlying variation may arise from multiple sources, not only the circadian pacemaker.

Our design decorrelates circadian phase from time awake. In the ultra-short sleep–wake protocol, frequent sleep opportunities were intended to reduce and stabilise homeostatic influences on visual thresholds. The present null result therefore suggests that previously reported time-of-day effects may be dominated by non-circadian factors or may arise in visual mechanisms not probed by our conedirected photopic temporal modulation.

Two processes could, in principle, explain the discrepancy between earlier conclusions, our initial hypotheses, and the present results. First, the absence of a detected rhythm in functional performance measures could reflect a phase shift or reset of the circadian clock induced by the intervention. The hormone data argue against that interpretation. Salivary melatonin profiles indicate intact central circadian signalling during the forced desynchrony protocol: the suprachiasmatic nucleus (SCN) controls melatonin release from the pineal gland via the sympathetic nervous system, and the resulting salivary dynamics were physiologically plausible throughout the protocol. Circadian periods were also comparable to those reported in the literature, with a mean duration slightly longer than 24 hours (Duffy et al., 2011). Although participants necessarily received intermittent light exposure while performing the visual tasks, the melatonin profiles do not suggest suppressive light effects (Gronfier, Wright, Kronauer, Jewett, & Czeisler, 2004). We cannot entirely exclude phase delays in melatonin onset, as those may be functionally independent (Rahman et al., 2018), but the fitted periods were well within the expected range (Duffy et al., 2011). Given the strength of evidence against circadian modulation in the behavioural data, it is unlikely that a phase delay in melatonin dissociated the relationship between salivary melatonin and visual performance.

Unlike other work that has focused on spatial contrast sensitivity or increment-threshold tasks (Andrade et al., 2019), we used TCS to compare stimulus directions that weight luminance and chromatic mechanisms differently while avoiding additional confounds from spatial optics. A spatial task would have introduced chromatic aberration for chromatic stimuli (Marimont & Wandell, 1994), and any day-night change in those optical properties would have been difficult to distinguish from neural or circadian effects. The temporal task therefore provided a cleaner, though narrower, test of whether cone-directed visual performance tracks circadian phase.

Descriptively, the task behaved as expected from prior work on temporal tuning (Gelfand & Horwitz, 2018; Dobkins et al., 1997). Sensitivity to L–M modulation was relatively higher at 2 Hz than at 8 Hz, whereas sensitivity to L+M+S modulation was relatively higher at 8 Hz than at 2 Hz. We interpret this pattern as a manipulation check showing that the task sampled different portions of the temporal response functions. At the same time, we do not claim that either frequency exclusively isolated one mechanism. The measured thresholds should instead be understood as being weighted differently toward chromatic and luminance contributions.

Taken together, the evidence indicates no detectable near-24-hour modulation of the specific behavioural measure tested here, namely cone-directed temporal contrast sensitivity under controlled photopic viewing conditions. This does not imply that all visual functions are circadianly invariant. Rather, it narrows the inference: previously reported time-of-day effects on visual performance may depend on sleep pressure, vigilance, posture, feeding state, pupil-linked changes in retinal illuminance, rod or melanopsin contributions, or on visual tasks that probe mechanisms beyond those sampled here.

Attempts to reconcile retinal rhythmicity with null behavioural effects remain possible. One candidate is that rhythmic modulation acts more strongly in rods than in cones. Hankins et al. (1998) reported diurnal variation in ERG components that may depend on rod-mediated processes, and human studies have suggested larger time-of-day effects under scotopic or mesopic conditions than under clearly photopic ones (O’Keefe & Baker, 1987; Tassi & Pins, 1997). Another possibility is that local retinal clocks and the central pacemaker are not phase-aligned (Roenneberg et al., 1992). Our data cannot directly arbitrate between these explanations, but they do indicate that any such processes did not produce a detectable circadian signature in the cone-directed temporal thresholds measured here.

### Limitations

#### Reliance on behavioural outcome measures

One limitation of the study is its reliance on a psychophysical task to explore circadian effects in post-receptoral mechanisms. This approach depends on behavioural responses and therefore remains susceptible to fluctuations in motivation and vigilance, even though the forced desynchrony design was chosen specifically to reduce large variations in sleep pressure. Future studies could complement psychophysical outcomes with electrophysiological or neurophysiological measures to test whether those modalities converge on the same conclusion. In particular, visual evoked potentials could provide a useful comparison with prior work while avoiding some response-related variance (Tsoneva, Garcia-Molina, & Desain, 2023).

#### Residual homeostatic and behavioural influences

The forced desynchrony protocol was designed primarily to reduce and stabilise sleep-pressure-related homeostatic influences. It does not guarantee invariance of all physiological systems across the protocol. Metabolic, autonomic, and other homeostatic processes may still have varied to some extent despite the regular schedule and controlled environment. Our inference is therefore narrower: under a protocol that strongly reduces major behavioural confounds, we found no convincing evidence that circadian phase alone modulated the temporal contrast sensitivity measure used here.

#### Scope of the psychophysical measure

The study used one psychophysical outcome measure with stimuli presented at two symmetric parafoveal locations. This design prioritised a repeatable, efficient assay that could be embedded in the circadian protocol, but it also limits generalisation. The task is sensitive to relative contrast thresholds, not to every possible change in absolute responsivity. If circadian modulation acted primarily by changing absolute cone gain while leaving relative contrast relationships intact, the present task could underestimate such an effect. Likewise, any stable left-right sensitivity asymmetry within an observer would contribute variance rather than a circadian signal.

#### Artificial pupil and ecological validity

The 1.5 mm artificial pupil was used to hold retinal irradiance constant despite endogenous pupil fluctuations. This was important for isolating changes in sensitivity from changes in retinal illuminance, but it also reduces ecological validity because natural viewing involves a closed loop between pupil size and retinal irradiance. We therefore cannot exclude the possibility that pupil-mediated time-of-day effects contribute to visual performance under ordinary viewing conditions even though they were intentionally suppressed here.

#### Nominal rather than observer-specific silent substitution

Our stimuli were generated using standard-observer cone fundamentals and a three-primary display. This provides principled control of the targeted stimulus directions, but it does not guarantee perfect isolation in every participant because observer-specific L:M cone ratio, prereceptoral filtering, and macular pigment density were not measured. The results should therefore be interpreted as applying to nominally cone-directed stimulus directions rather than to perfectly purified post-receptoral mechanisms.

#### Other photoreceptor classes

**\**Finally, we did not explicitly silence rods or melanopsin. Recent studies demonstrate that rods can contribute under photopic conditions (Tikidji-Hamburyan et al., 2017), and melanopsin-mediated signals can influence temporal and visual detection tasks (Zele, Feigl, Adhikari, Maynard, & Cao, 2018; Uprety, Adhikari, Feigl, & Zele, 2022). With a three-primary display it is not possible to control all relevant photoreceptor classes simultaneously. Future work on circadian rhythmicity of visual sensitivity should therefore test rod- and melanopsin-related contributions more directly, ideally with multi-primary stimulation devices and with scotopic, mesopic, and photopic viewing conditions.

### Strengths

The study has three main strengths. Foremost, it combines a state-of-the-art circadian protocol designed to separate circadian phase from major sleep–wake-related confounds with targeted cone-directed stimulation using silent substitution.

#### Protocol to disentangle circadian and homeostatic regulation effects

The forced desynchrony protocol with an ultra-short sleep– wake cycle is a central strength of the design. By deliberately misaligning behavioural rhythms (e.g. sleep and wake times) with endogenous circadian phase, the protocol greatly reduces the interpretational ambiguity that affects conventional time-of-day studies. The present null finding is therefore more informative than a null result obtained under habitual daily schedules.

#### Mechanism-targeted stimulation

Compared with monochromatic or broadly filtered stimuli, silent substitution offers stronger control over the cone-contrast directions entering the visual system (Spitschan & Woelders, 2018). Even though this control is standard-observer based rather than observer-specific, it allowed us to compare nominal luminance, L–M, and S-cone-directed modulations within a single coherent framework.

#### Criterion-free task

The use of a 2AFC task is also advantageous in this context. Relative to yes-no detection tasks, 2AFC reduces the influence of individual response criteria and allows thresholds to be estimated from a staircase driven by hits and misses rather than by subjective “seen/unseen” reports.

## Conclusions

In this study, we decoupled circadian phase from major sleep–wake-related influences over 40 hours and measured cone-directed temporal contrast sensitivity in humans to probe post-receptoral processing of luminance and chromatic stimulus directions across the circadian cycle. We found no evidence for a 24-hour modulation of human temporal contrast sensitivity. Bayes factor model comparisons decisively favoured a null model with participant-specific intercepts over models including oscillatory components; estimated amplitudes were near zero and phases were dispersed across participants. Within these highly controlled conditions, any SCN-linked circadian effects on cone-directed temporal contrast sensitivity, if present, appear negligible. This suggests that previously reported diurnal variations in visual performance may primarily reflect homeostatic factors, pupil-linked changes in retinal illuminance, or contributions from non-cone pathways rather than a robust circadian modulation of the measure studied here. Generalisation to other visual regimes (for example, mesopic or scotopic conditions, spatial contrast sensitivity, or tasks with observer-specific optical variation) remains to be established. Our findings therefore underscore both the value of rigorous circadian protocols and the importance of limiting conclusions to the specific visual measure and viewing conditions that were actually tested.

## Acknowledgements

We gratefully acknowledge the mechanical workshop of the Max Planck Institute for Biological Cybernetics for their crucial contributions in designing, and setting up a sleep laboratory for the purpose of this study. We thank Katharina Stein, and Constanza Bravo for their efforts during the preparation of the study and during piloting the experimental procedure, which provided valuable insights for refining the study design. We also extend our appreciation to our colleague Salma M. Thalji and the team of student assistants, Lisa Gagelmann and Emin Kılıç for their dedicated work during data collection.

## APPENDIX A Participant-specific fit for salivary melatonin concentration

Participant-level fits of the periodic model to the salivary melatonin concentrations are shown in Figure A1, and the corresponding identification of the participant-specific period *τ* from the goodness-of-fit search is shown in Figure A2.

Calculations to obtain an estimate for participant-specific period *τ*_*j*_ were performed under various assumptions. First, the melatonin period reflects the current rather than the endogenous circadian rhythm. The circadian pacemaker was subjected to entrainment before entering the laboratory protocol. In order for the circadian rhythm to be “free running”, i.e. for the salivary melatonin and other indicators to operate on the endogenous period, multiple days up to a few weeks in dim light are necessary. A dim light period of 13 hours (4 hours of the dim-light evening protocol, 8 hours of adaptation night, and 1 hour of setup after wake-up) preceding the forced desynchrony protocol possibly induces a drift from the entrained state but is likely not sufficient to establish a “free-running” rhythm. Second, the assumption that the total experimental light exposure delivered during the vision tests does not lead to a net phase delay or advance throughout the experiment, i.e. period and phase does not change from one cycle to the next. To summarise, the identified *τ*_*j*_ for an individual *j* is expected to reflect the participant’s *current* circadian period during the laboratory stay rather than the endogenous free-running period of the circadian pacemaker.

**Figure A1.**
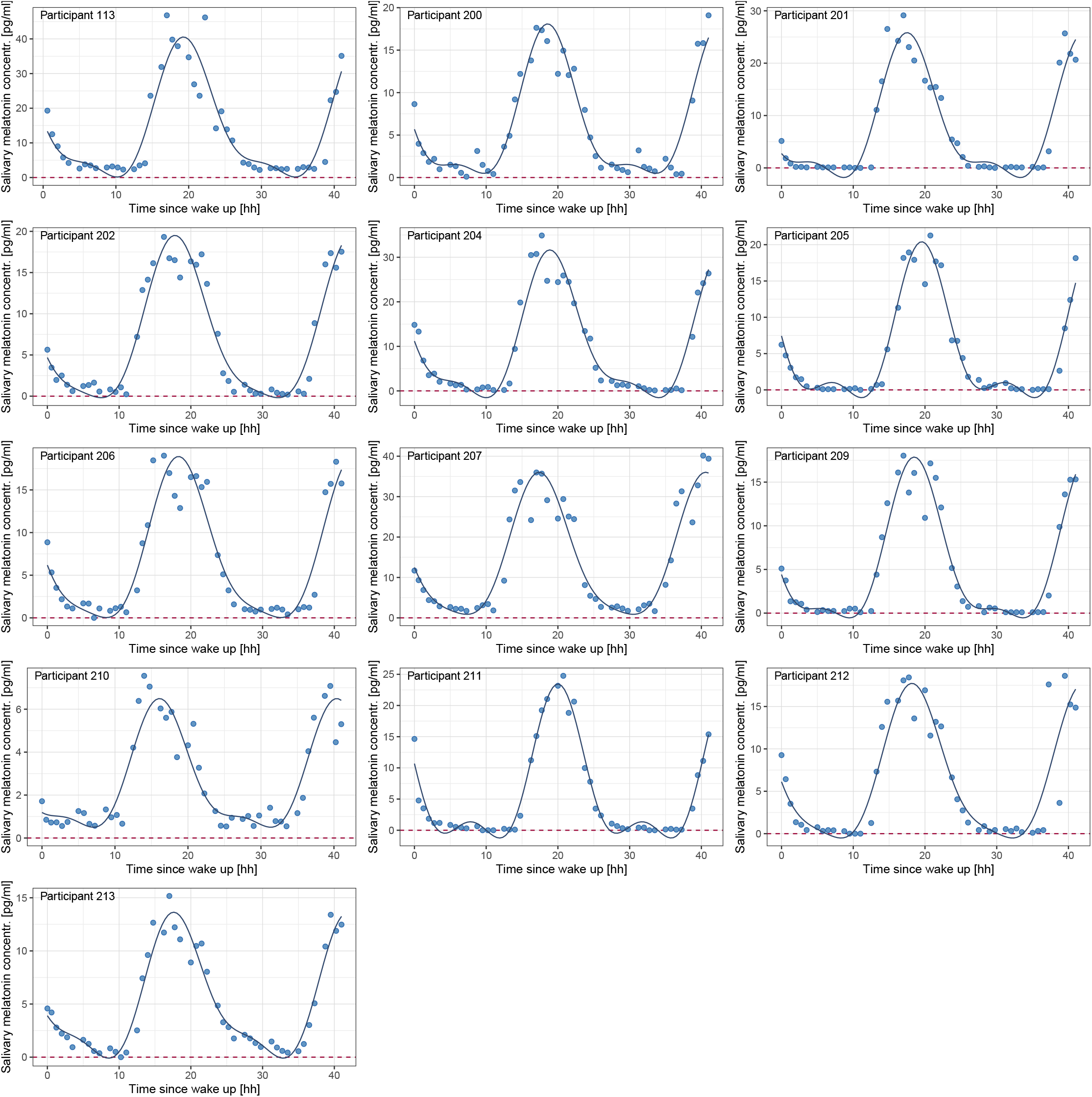
Participant-level salivary melatonin concentrations (pg/mL) as a function of time since wake-up. Blue circles are measurements for each participant; the solid curve is the periodic model fit comprising the fundamental and second-harmonic components. The horizontal red dashed line marks zero concentration. Each panel shows an individual participant’s data and corresponding fit.

**Figure A2.**
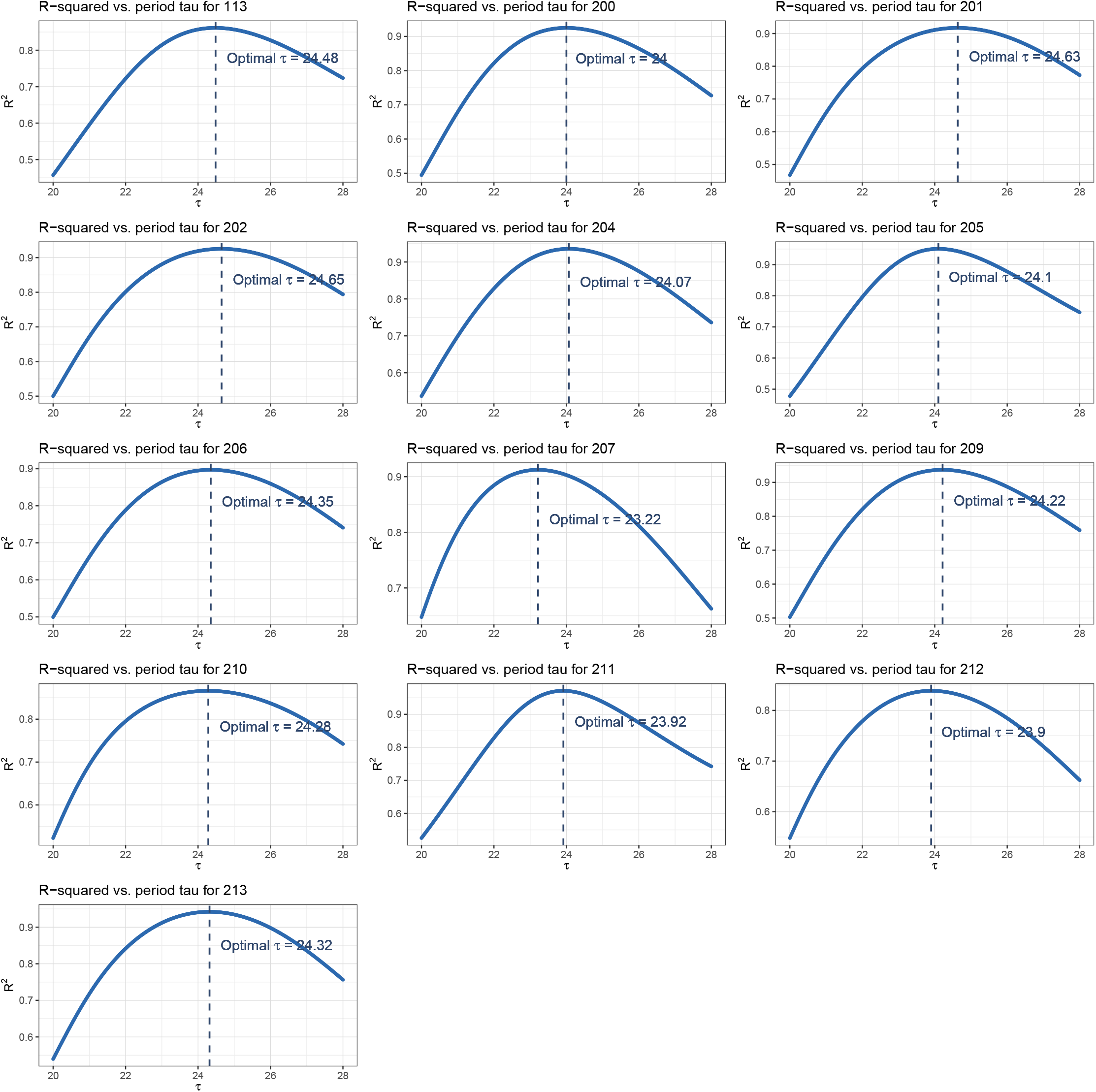
Identification of the participant-specific period *τ* of the periodic secretion of melatonin, as measured in saliva concentration. For each participant, the same bi-harmonic model was refit across a range of plausible *τ* values, from 20 to 28 h in 1-minute increments, and the coefficient of determination (R^2^) is plotted as a function of *τ* . The vertical dashed line marks the *τ* that maximizes R^2^, indicating the optimal period for that participant.

## APPENDIX B Model results from fitting temporal contrast sensitivity

The supplementary models were fit to the unlogged outcome variable. Their results were substantively unchanged and led to the same conclusions. Model statistics are reported in Tables B1 (coefficients) and B2 (fit statistics), and all ln BF estimates fell within [−18.941, −9.41].

**Table B1:** Fixed- and random-effect estimates and derived group-level amplitude and phase for the supplementary models fit to the unlogged temporal contrast sensitivity. Within each model block, the left *Estimate*/*CI* columns refer to the full (circadian) model and the right columns to the null model. Fit statistics are reported separately in Table B2.

| Model | Predictor | Full model |  |  | Null model |  |  |
| --- | --- | --- | --- | --- | --- | --- | --- |
|  |  | Estimate | CI |  | Estimate | CI |  |
|  |  |  | 2.5% | 97.5% |  | 2.5% | 97.5% |
| L–M (red-green), 2 Hz | Fixed effects |  |  |  |  |  |  |
|  | Intercept | 1.178 | 0.900 | 1.453 | 1.191 | 0.921 | 1.457 |
| | $\beta$ | -0.023 | -0.153 | 0.109 | | | |
| | $\alpha$ | 0.075 | -0.051 | 0.202 | | | |
|  | Random effects |  |  |  |  |  |  |
|  | SD(Intercept) | 0.442 | 0.262 | 0.745 | 0.431 | 0.259 | 0.720 |
| | SD($\beta$) | 0.091 | 0.004 | 0.267 | | | |
| | SD($\alpha$) | 0.075 | 0.003 | 0.226 | | | |
| | $\rho(\text{Intercept}, \beta)$ | -0.056 | -0.870 | 0.827 | | | |
| | $\rho(\text{Intercept}, \alpha)$ | 0.076 | -0.834 | 0.886 | | | |
|  |  | 0.029 | -0.873 | 0.889 |  |  |  |
|  | Derived estimates |  |  |  |  |  |  |
|  | Amplitude | 0.079 |  |  |  |  |  |
|  | Phase | -0.296 |  |  |  |  |  |
| L–M (red-green), 8 Hz | Fixed effects |  |  |  |  |  |  |
|  | Intercept | 0.897 | 0.658 | 1.136 | 0.904 | 0.679 | 1.129 |
| | $\beta$ | 0.007 | -0.088 | 0.102 | | | |
| | $\alpha$ | 0.040 | -0.088 | 0.168 | | | |
|  | Random effects |  |  |  |  |  |  |
|  | SD(Intercept) | 0.387 | 0.235 | 0.648 | 0.366 | 0.224 | 0.606 |
| | SD($\beta$) | 0.067 | 0.003 | 0.193 | | | |
| | SD($\alpha$) | 0.147 | 0.012 | 0.328 | | | |
| | $\rho(\text{Intercept}, \beta)$ | 0.155 | -0.784 | 0.899 | | | |
| | $\rho(\text{Intercept}, \alpha)$ | -0.087 | -0.767 | 0.674 | | | |
|  |  | 0.147 | -0.810 | 0.911 |  |  |  |
|  | Derived estimates |  |  |  |  |  |  |
|  | Amplitude | 0.040 |  |  |  |  |  |
|  | Phase | 0.178 |  |  |  |  |  |
| L+M+S (luminance), 2 Hz | Fixed effects |  |  |  |  |  |  |

**Table B1 – continued from previous page**
| *Model* | *Predictor* | *Full model* |  |  | *Null model* |  |  |
| --- | --- | --- | --- | --- | --- | --- | --- |
|  |  | *Estimate* | *CI* |  | *Estimate* | *CI* |  |
|  |  |  | *2.5%* | *97.5%* |  | *2.5%* | *97.5%* |
|  | Intercept | 0.176 | 0.148 | 0.203 | 0.177 | 0.149 | 0.205 |
| | $\beta$ | -0.005 | -0.028 | 0.019 | | | |
| | $\alpha$ | 0.009 | -0.017 | 0.035 | | | |
|  | *Random effects* |  |  |  |  |  |  |
|  | SD(Intercept) | 0.039 | 0.018 | 0.071 | 0.041 | 0.020 | 0.074 |
| | SD($\beta$) | 0.021 | 0.001 | 0.052 | | | |
| | SD($\alpha$) | 0.025 | 0.001 | 0.061 | | | |
| | $\rho$(Intercept, $\beta$) | -0.430 | -0.958 | 0.575 | | | |
| | $\rho$(Intercept, $\alpha$) | 0.290 | -0.625 | 0.920 | | | |
|  |  | -0.180 | -0.914 | 0.773 |  |  |  |
|  | *Derived estimates* |  |  |  |  |  |  |
|  | Amplitude | 0.010 |  |  |  |  |  |
|  | Phase | -0.507 |  |  |  |  |  |
| L+M+S (luminance), 8 Hz | *Fixed effects* |  |  |  |  |  |  |
|  | Intercept | 0.404 | 0.338 | 0.470 | 0.411 | 0.347 | 0.476 |
| | $\beta$ | -0.016 | -0.074 | 0.042 | | | |
| | $\alpha$ | 0.042 | -0.014 | 0.098 | | | |
|  | *Random effects* |  |  |  |  |  |  |
|  | SD(Intercept) | 0.091 | 0.035 | 0.172 | 0.090 | 0.036 | 0.167 |
| | SD($\beta$) | 0.049 | 0.002 | 0.133 | | | |
| | SD($\alpha$) | 0.040 | 0.002 | 0.114 | | | |
| | $\rho$(Intercept, $\beta$) | 0.112 | -0.793 | 0.878 | | | |
| | $\rho$(Intercept, $\alpha$) | 0.145 | -0.794 | 0.905 | | | |
|  |  | -0.089 | -0.905 | 0.849 |  |  |  |
|  | *Derived estimates* |  |  |  |  |  |  |
|  | Amplitude | 0.045 |  |  |  |  |  |
|  | Phase | -0.359 |  |  |  |  |  |
| S (blue-yellow), 2 Hz | *Fixed effects* |  |  |  |  |  |  |
|  | Intercept | 0.173 | 0.134 | 0.212 | 0.175 | 0.137 | 0.212 |
| | $\beta$ | -0.009 | -0.027 | 0.009 | | | |
| | $\alpha$ | 0.007 | -0.011 | 0.024 | | | |
|  | *Random effects* |  |  |  |  |  |  |
|  | SD(Intercept) | 0.063 | 0.037 | 0.106 | 0.061 | 0.037 | 0.102 |
| | SD($\beta$) | 0.014 | 0.001 | 0.039 | | | |
| | SD($\alpha$) | 0.011 | 0.000 | 0.032 | | | |
| | $\rho$(Intercept, $\beta$) | -0.132 | -0.889 | 0.784 | | | |
| | $\rho$(Intercept, $\alpha$) | 0.062 | -0.838 | 0.883 | | | |
|  |  | -0.058 | -0.898 | 0.864 |  |  |  |

**Table B1 – continued from previous page**
| Model | Predictor | Full model |  |  | Null model |  |  |
| --- | --- | --- | --- | --- | --- | --- | --- |
|  |  | Estimate | CI |  | Estimate | CI |  |
|  |  |  | 2.5% | 97.5% |  | 2.5% | 97.5% |
|  | Derived estimates |  |  |  |  |  |  |
|  | Amplitude | 0.011 |  |  |  |  |  |
|  | Phase | -0.936 |  |  |  |  |  |
| S (blue-yellow), 8 Hz | Fixed effects |  |  |  |  |  |  |
|  | Intercept | 0.097 | 0.069 | 0.126 | 0.098 | 0.070 | 0.126 |
| | $\beta$ | 0.004 | -0.003 | 0.012 | | | |
| | $\alpha$ | 0.006 | -0.002 | 0.014 | | | |
|  | Random effects |  |  |  |  |  |  |
|  | SD(Intercept) | 0.047 | 0.030 | 0.078 | 0.047 | 0.030 | 0.077 |
| | SD($\beta$) | 0.005 | 0.000 | 0.014 | | | |
| | SD($\alpha$) | 0.007 | 0.000 | 0.017 | | | |
| | $\rho(\text{Intercept}, \beta)$ | 0.113 | -0.817 | 0.898 | | | |
| | $\rho(\text{Intercept}, \alpha)$ | 0.342 | -0.654 | 0.940 | | | |
|  |  | 0.082 | -0.848 | 0.901 |  |  |  |
|  | Derived estimates |  |  |  |  |  |  |
|  | Amplitude | 0.007 |  |  |  |  |  |
|  | Phase | 0.605 |  |  |  |  |  |

**Table B2:**
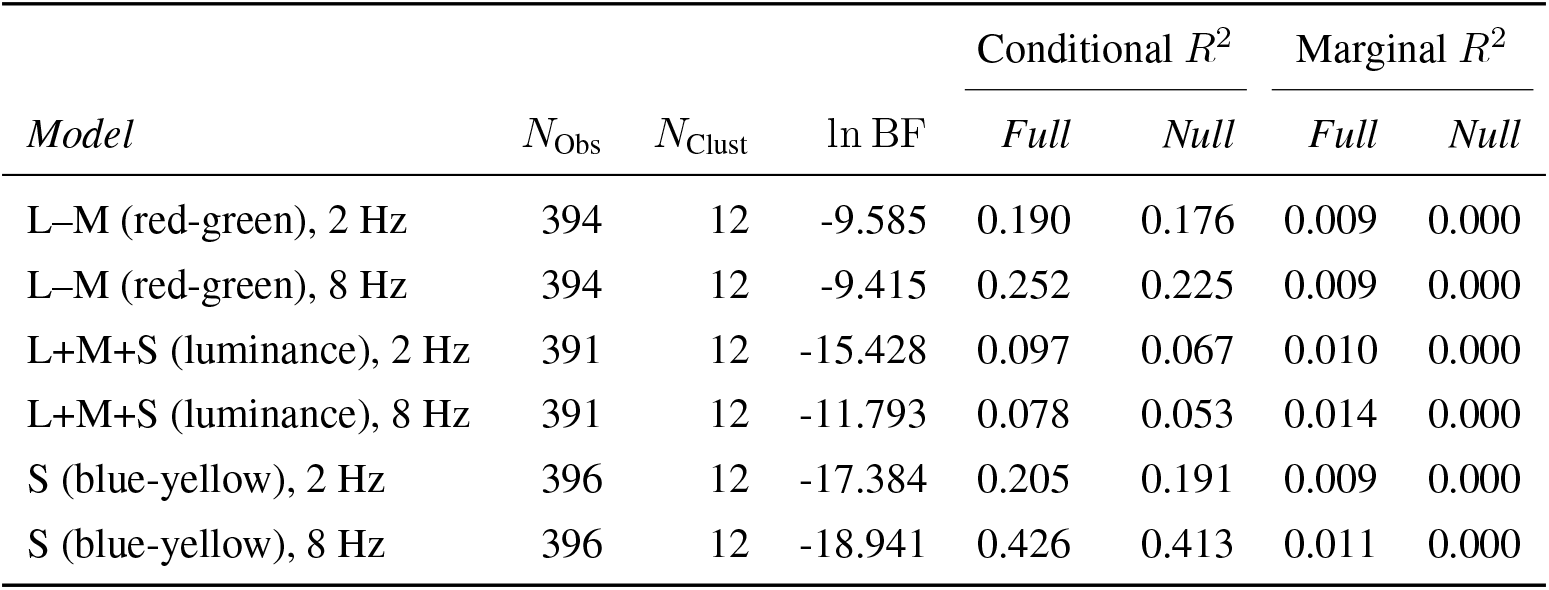
Sample sizes, model-comparison evidence and explained variance for the supplementary models fit to the unlogged temporal contrast sensitivity. Conditional and marginal *R*^2^ are reported for the full (circadian) and null models. Coefficient estimates are reported in Table B1.

| *Model* | $N_{\text{Obs}}$ | $N_{\text{Clust}}$ | $\ln \text{BF}$ | *Conditional $R^2$* | | *Marginal $R^2$* | |
| --- | --- | --- | --- | --- | --- | --- | --- |
|  |  |  |  | *Full* | *Null* | *Full* | *Null* |
| L–M (red-green), 2 Hz | 394 | 12 | -9.585 | 0.190 | 0.176 | 0.009 | 0.000 |
| L–M (red-green), 8 Hz | 394 | 12 | -9.415 | 0.252 | 0.225 | 0.009 | 0.000 |
| L+M+S (luminance), 2 Hz | 391 | 12 | -15.428 | 0.097 | 0.067 | 0.010 | 0.000 |
| L+M+S (luminance), 8 Hz | 391 | 12 | -11.793 | 0.078 | 0.053 | 0.014 | 0.000 |
| S (blue-yellow), 2 Hz | 396 | 12 | -17.384 | 0.205 | 0.191 | 0.009 | 0.000 |
| S (blue-yellow), 8 Hz | 396 | 12 | -18.941 | 0.426 | 0.413 | 0.011 | 0.000 |

## APPENDIX C Diurnal control experiment

We re-invited two participants in July and August 2025 to conduct control experiments diurnally and with different viewing conditions. Aim of this control experiment was 1. to compare contrast sensitivity estimates obtained with our experimental setup employed during the forced desynchrony with the pupil diameter being constrained by an external pupil (“constrained-pupil”), to those obtained with less controlled viewing conditions (“free-viewing”). In the “free-viewing” condition, participants viewed stimuli on monitor without any devices restricting view of their dominant eye. The “constrained-pupil” condition replicated the experimental conditions of the main study, i.e. monocular viewing with the dominant eye through a pupil relay system which restricts the effective pupil diameter to 1.5 mm, the non-dominant eye was always covered with a one-sided eye patch; eye dominance was determined using the Miles test as described for the main study. 2. to report empirical pupil variability during the experimental procedure and across the day; 3. and finally, to explore the possibility of diurnal pattern rather than circadian modulation of the psychophysical performance measure. Control experiment conducted diurnally from 8 AM to 8:30 PM, it consisted of 10 blocks lasting 1:15 hours each, i.e. a third of the duration of one block during the forced desynchrony protocol of the main study. We began each session with the “constrained-pupil” condition and subsequently alternated between “constrained-pupil” and “free-viewing” conditions. In both conditions, the effective distance between the stimulus display and the participant’s pupil was held constant at 1.5 m. In the “constrained-pupil” condition, an external pupil aperture was positioned 1.5 m from the participant. A pupil relay system projected the aperture over a 15 cm optical path onto the participant’s pupil without net magnification. Pupil size was recorded independently of the psychophysical responses using Pupil Core (Pupil Labs GmbH). Each recording included (i) a standardised calibration procedure performed before every experimental run and(ii) pupil dynamics during stimulus presentation (stimuli of varying contrast at two temporal frequencies, within one condition), plus a short period after the run ended. During calibration, participants fixated straight ahead, then shifted gaze (without moving the head) to instructed directions (left, up, right, down), returning to look straight ahead in the end. Pupil recordings were matched to the threshold estimate from the corresponding experimental run. Pupil diameter was estimated in millimetres using Pupil Core’s 3D eye model derived from the calibration procedure. Initial preprocessing was performed in Pupil Core v3.5 (open-source) using default settings, including automated blink detection, and the resulting pupil-diameter time series were exported for analysis. Post-processing encompassed a plausibility filter, which excluded pupil sizes ]0.5; 10[ mm which were less than 2% across all runs; and a high-confidence filter, allowing only samples a confidence score > 0.8 in the reported Pupil Labs detection.

We obtained summary statistics of the pupil diameter of each recording; results are visualised in Fig. C2. Summary statistics were derived from the test-specific summary statistics and reported in Table C2, and more detailed with per-block summary statistics in Table C3. The contrast sensitivities from the diurnal experiment are reported and visualised in Tables C1, and visualised in Fig. C1. There is a marked difference between the “free-viewing” and the “constrained-pupil” condition.

**Table C1:** Summary statistics of contrast sensitivity per participant, condition, temporal frequency, and context (mean with std. dev. in parentheses). Values reflect all trials pooled across experimental blocks.

| Participant ID | Condition | Temporal frequency [Hz] | Constrained pupil | Free viewing |
| --- | --- | --- | --- | --- |
| 113 | L+M+S (luminance) | 2 | 0.154 (0.071) | 0.173 (0.038) |
|  |  | 8 | 0.430 (0.143) | 0.439 (0.120) |
|  | L–M (red-green) | 2 | 0.796 (0.254) | 1.958 (0.409) |
|  |  | 8 | 0.719 (0.277) | 1.817 (0.598) |
|  | S (blue-yellow) | 2 | 0.271 (0.553) | 0.244 (0.096) |
|  |  | 8 | 0.090 (0.056) | 0.243 (0.245) |
| 202 | L+M+S (luminance) | 2 | 0.208 (0.053) | 0.239 (0.067) |
|  |  | 8 | 0.471 (0.144) | 0.640 (0.192) |
|  | L–M (red-green) | 2 | 1.101 (0.213) | 3.438 (1.652) |
|  |  | 8 | 0.996 (0.190) | 2.518 (0.698) |
|  | S (blue-yellow) | 2 | 0.196 (0.073) | 0.405 (0.311) |
|  |  | 8 | 0.123 (0.026) | 0.256 (0.054) |

**Figure C1.**
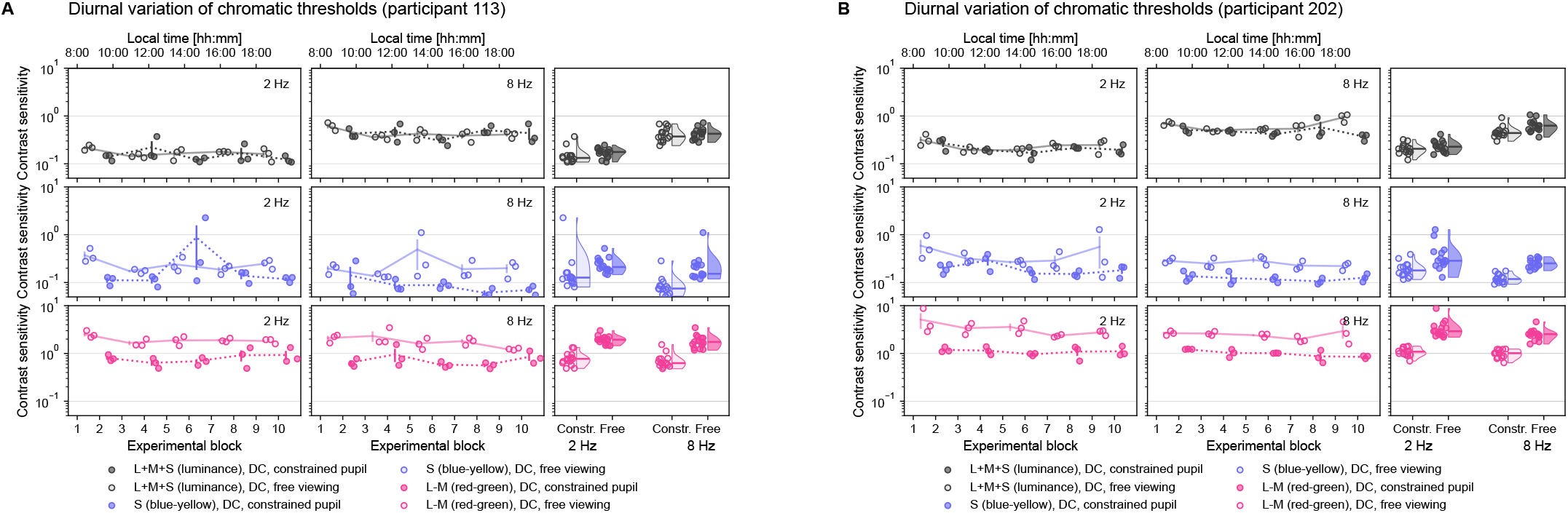
Diurnal dynamics in temporal contrast sensitivity to different stimulus conditions (rows; “L+M+S (luminance)”, “S (blue-yellow)” or “L–M (red-green)”), and modulation frequencies (first two columns for 2 Hz and 8 Hz); last column contains raincloud plots with marginal distributions of contrast sensitivity estimates and distributions.

**Table C2:** Summary statistics of pupil diameter per participant and condition (mean with std. dev. in parentheses, in mm). Values are computed from free-viewing blocks only, pooled across all within-block recordings.

| Participant ID | Condition | Pupil diameter [mm] |
| --- | --- | --- |
| 113 | L+M+S (luminance) | 4.228 (0.615) |
|  | L–M (red-green) | 4.568 (0.619) |
|  | S (blue-yellow) | 4.465 (0.743) |
| 202 | L+M+S (luminance) | 5.034 (0.531) |
|  | L–M (red-green) | 5.087 (0.660) |
|  | S (blue-yellow) | 5.135 (0.497) |

**Figure C2.**
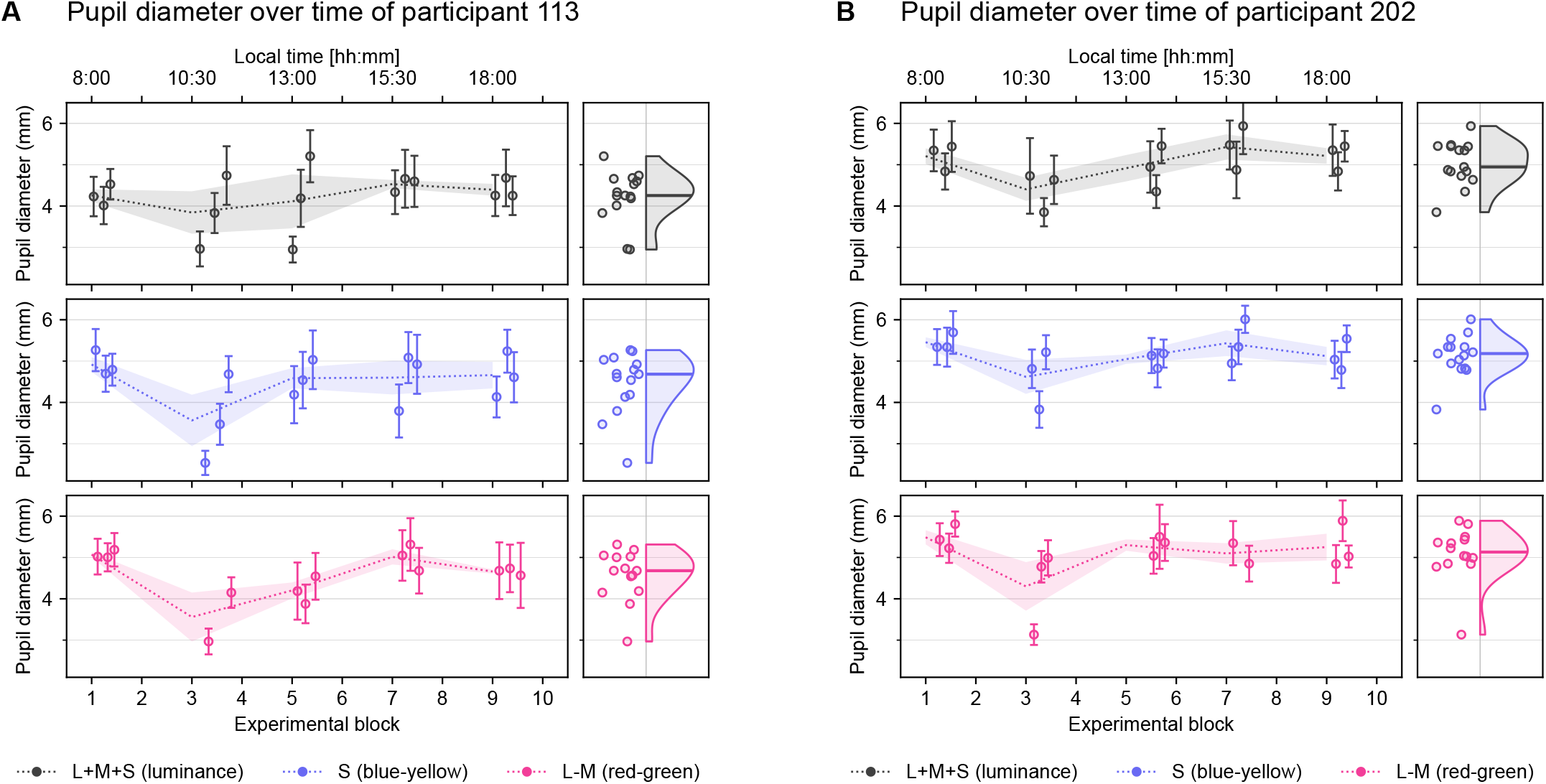
Diurnal dynamics of pupil diameter in different experimental stimulus conditions (“L+M+S (luminance)”, “S (blue-yellow)” or “L–M (red-green)”) every other block. A and B show data of different participants with two panels each. Left panels show mean pupil diameter and standard deviation of individual runs as scatterplot, and dotted line to connects per-block means with ribbon indicating standard mean error. Right panels include raincloud plots of marginal distributions of mean pupil diameter across the day (8:00 to 20:30 local time) for the respective participants. Recordings were obtained under monocular viewing conditions from dominant eye only; non-dominant eye was covered by one-sided eye patch.

**Table C3:** Summary statistics of mean pupil diameter across conditions (one row per participant and block).

| Condition | Participant ID | Block | Pupil diameter [mm] |  |  |  |
| --- | --- | --- | --- | --- | --- | --- |
|  |  |  | *Mean* | *Std. dev.* | *Min* | *Max* |
| L+M+S (luminance) | 113 | 0 | 4.257 | 0.258 | 0.794 | 5.307 |
|  |  | 2 | 3.844 | 0.888 | 0.728 | 6.062 |
|  |  | 4 | 4.113 | 1.131 | 0.764 | 6.488 |
|  |  | 6 | 4.531 | 0.172 | 0.793 | 7.340 |
|  |  | 8 | 4.395 | 0.247 | 0.529 | 6.204 |
|  | 202 | 0 | 5.207 | 0.324 | 0.725 | 6.353 |
|  |  | 2 | 4.405 | 0.482 | 0.548 | 5.831 |
|  |  | 4 | 4.915 | 0.552 | 0.724 | 6.077 |
|  |  | 6 | 5.428 | 0.532 | 0.960 | 7.039 |
|  |  | 8 | 5.212 | 0.327 | 0.907 | 6.099 |
| S (blue-yellow) | 113 | 0 | 4.917 | 0.306 | 0.934 | 6.144 |
|  |  | 2 | 3.563 | 1.077 | 0.532 | 5.612 |
|  |  | 4 | 4.586 | 0.424 | 0.854 | 8.417 |
|  |  | 6 | 4.600 | 0.704 | 0.776 | 6.455 |
|  |  | 8 | 4.660 | 0.555 | 0.945 | 6.929 |
|  | 202 | 0 | 5.457 | 0.204 | 0.883 | 6.312 |
|  |  | 2 | 4.618 | 0.713 | 0.736 | 5.936 |
|  |  | 4 | 5.046 | 0.194 | 0.610 | 5.864 |
|  |  | 6 | 5.432 | 0.539 | 0.883 | 6.492 |
|  |  | 8 | 5.121 | 0.385 | 0.824 | 6.048 |
| L-M (red-green) | 113 | 0 | 5.070 | 0.101 | 1.000 | 5.967 |
|  |  | 2 | 3.559 | 0.837 | 0.603 | 4.948 |
|  |  | 4 | 4.202 | 0.334 | 0.783 | 6.196 |
|  |  | 6 | 5.014 | 0.318 | 0.884 | 9.592 |
|  |  | 8 | 4.660 | 0.086 | 0.529 | 9.918 |
|  | 202 | 0 | 5.488 | 0.296 | 0.913 | 6.256 |
|  |  | 2 | 4.300 | 1.016 | 0.586 | 5.489 |
|  |  | 4 | 5.300 | 0.237 | 0.735 | 6.710 |
|  |  | 6 | 5.098 | 0.350 | 0.804 | 6.091 |
|  |  | 8 | 5.251 | 0.558 | 0.926 | 6.551 |

